# Targeting the pathological network: feasibility of network-based optimization of transcranial magnetic stimulation coil placement for treatment of psychiatric disorders

**DOI:** 10.1101/2022.10.23.513193

**Authors:** Zhengcao Cao, Xiang Xiao, Yang Zhao, Yihan Jiang, Cong Xie, Marie-Laure Paillère-Martinot, Eric Artiges, Zheng Li, Zafiris J. Daskalakis, Yihong Yang, Chaozhe Zhu

## Abstract

It has been recognized that the efficacy of TMS-based modulation may depend on the network profile of the stimulated regions throughout the brain. However, what profile of this stimulation network optimally benefits treatment outcomes is yet to be addressed. The answer to the question is crucial for informing network-based optimization of stimulation parameters, such as coil placement, in TMS treatments. In this study, we aimed to investigate the feasibility of taking a disease-specific network as the target of stimulation network for guiding individualized coil placement in TMS treatments. We present here a novel network-based model for TMS targeting of the pathological network. First, combining E-field modeling and resting-state functional connectivity, stimulation networks were modeled from locations and orientations of the TMS coil. Second, the spatial anti-correlation between the stimulation network and the pathological network of a given disease was hypothesized to predict the treatment outcome. The proposed model was validated to predict treatment efficacy from the position and orientation of TMS coils in two depression cohorts and one auditory verbal hallucinations cohort. We further demonstrate the utility of the proposed model in guiding individualized TMS treatment for psychiatric disorders. In this proof-of-concept study, we demonstrated the feasibility of the novel network-based targeting strategy that uses the whole-brain, system-level abnormity of a specific psychiatric disease as a target. Results based on empirical data suggest that the strategy may potentially be utilized to identify individualized coil parameters for maximal therapeutic effects.

**Highlights:**

- Proposed a model of targeting pathological brain networks for pre-treatment TMS coil placement planning in the treatment of psychiatric disorders;
- Validated the network targeting model in three cohorts of patients with depression or auditory verbal hallucinations, via prediction of individual TMS treatment efficacy from the parameters of coil placement;
- Demonstrated the utility of the network targeting model in guiding individualized TMS coil placement.

## 1. Introduction

Transcranial magnetic stimulation (TMS) is a noninvasive neuromodulation technology that can modulate neural activity with high spatial sensitivity (Barker et al., 1985). Accumulating evidence has shown its potential as a clinical therapy for many psychiatric disorders (Rossini et al., 2010; Lefaucheur et al., 2014; Sale et al., 2015). However, the large variation in treatment efficacy across diseases and individual patients underscores the importance to improve the current TMS treatment protocols.

In TMS-based treatment, a major methodological issue is how to achieve optimal efficacy by choosing the parameters, particularly the position and orientation of the TMS coil (Fitzgerald, 2021). Traditionally, TMS coils are placed according to anatomically defined regions, e.g., dorsolateral prefrontal cortex (DLPFC) for major depressive disorder (MDD). TMS coils are usually placed on a specific site, e.g., 5-cm from the motor hotspot (George et al., 1994; Pascual-Leone et al., 1996), referring to scalp landmarks of the EEG 10-20 system (Herwig et al., 2003; Beam et al., 2009), or projecting to brain coordinates via a neuronavigation system (Herwig et al., 2001; Fitzgerald et al., 2009). However, the location of region-of-interest (ROI) alone is insufficient for guiding the optimal setting of TMS coils. First, within the targeted ROI, the distribution of the E-field generated by TMS further depends on the pose of the TMS coil relative to the gyrification of cortex underneath (Richter et al., 2013; Gomez-Tames et al., 2018). Accordingly, it is necessary to consider the location-and-orientation interaction when placing TMS coils for optimal outcomes, even in the case of motor-evoked potentials (Reijonen et al., 2020). Second, the treatment response of TMS may further depend on the specific functional network associated with cortical regions directly affected by the stimulation. TMS is capable of generating effects in remote brain regions connected to the local stimulating site (Bestmann et al., 2008; Eldaief et al., 2011; Reithler et al., 2011; Tik et al., 2017). Effective treatments are found to be accompanied by stimulation-induced changes in brain activity that occur in the downstream regions or their functional connectivity with the local region (Wang et al., 2014; Cash et al., 2019; Howard et al., 2020). Therefore, even when a given ROI is targeted, distinct functional networks can be affected by TMS in different individuals, and such variation of stimulation networks may account for the heterogeneity of the treatment response (Opitz et al., 2016; Cardenas et al., 2022). Resolving how the stimulation network mediates the relationship between the coil settings and the treatment outcome is critical for guiding the individualized optimization of TMS parameters.

For modeling the whole brain profile of the stimulation network from coil settings on an individual’s scalp, a previous work by Opitz et al (Opitz et al., 2016) described a general framework integrating the realistic E-field modeling (Windhoff et al., 2013) and resting-state functional connectivity (rsFC) mapping (Fox and Raichle, 2007; Fox et al., 2012, 2014a). This framework allows one to address TMS targeting at the network level. In a healthy cohort, this framework demonstrated how the stimulation networks vary among individuals when DLPFC was selected for treating MDD. However, it remains unclear what stimulation network profile will optimally benefit the clinical/behavioral outcome, which is crucial in guiding treatment for psychiatric disorders.

For determining beneficial stimulation network profiles, a “pathological network” of a specific psychiatric disease (e.g., the difference in brain activity between patients and controls) may serve as a potential target. Psychiatric disorders have been recognized as network disruptions (Silbersweig et al., 1995; Mayberg, 1997; Fornito and Bullmore, 2015; Braun et al., 2018). In MDD, multiple cortical and limbic nodes showing abnormal activity compared to healthy controls have been recognized to underpin the disease. Seminal research in depression has found that stimulation sites with stronger negative functional connectivity to the subgenual cingulate cortex (SGC), one deep node of the putative frontal-limbic network of depression, bear better treatment outcomes (Fox et al., 2012; Weigand et al., 2018). These findings inspire a hypothesis that the association between the stimulation network and the pathological network of a given disease may mediate the outcome drawn by TMS.

Based on this hypothesis, we propose a novel network targeting model for guiding individualized coil settings in treating psychiatric disorders. We first validated the feasibility of the proposed model in predicting treatment efficacy from TMS coil settings on individual scalps retrospectively on two cohorts of depression. Then, we further validated the feasibility to generalize this model to another disease, auditory verbal hallucinations (AVH). Finally, we demonstrated that optimized coil placement parameters vary between individual patients, which emphasizes the importance of individualized coil placement in TMS-based treatment.

## 2. Materials and Methods

### 2.1. Description of the Network Targeting Model

#### 2.1.1. Rationale of the model

The proposed model is based on the relationship between two conceptional networks: the stimulation network and the pathological network of a given disease. In the current scope, TMS parameters are limited to the position and orientation of TMS coil, and treatment outcome is defined by the change of disease severity measured with clinical scales. For a given setting of TMS coil parameters (Figure 1Ai), the TMS stimulation region is defined as the cortical region that is directly modulated by TMS, and estimated from FEM models based on the individual’s structural MRIs (Figure 1Aii). Then the stimulation network, defined as the profile of the whole-brain rsFC seeded from the stimulation region, was estimated from the voxel-wise connectome averaged from a large sample healthy cohort (Figures 1Aiii,Aiv). Individuals showing spatial anti-correlations between their stimulation networks (Figure 1B) and the pathological network of a given disease (Figure 1C) are hypothesized to be associated with effective treatment by TMS (Fox et al., 2014a)(Figure 1D).

**FIGURE 1.**
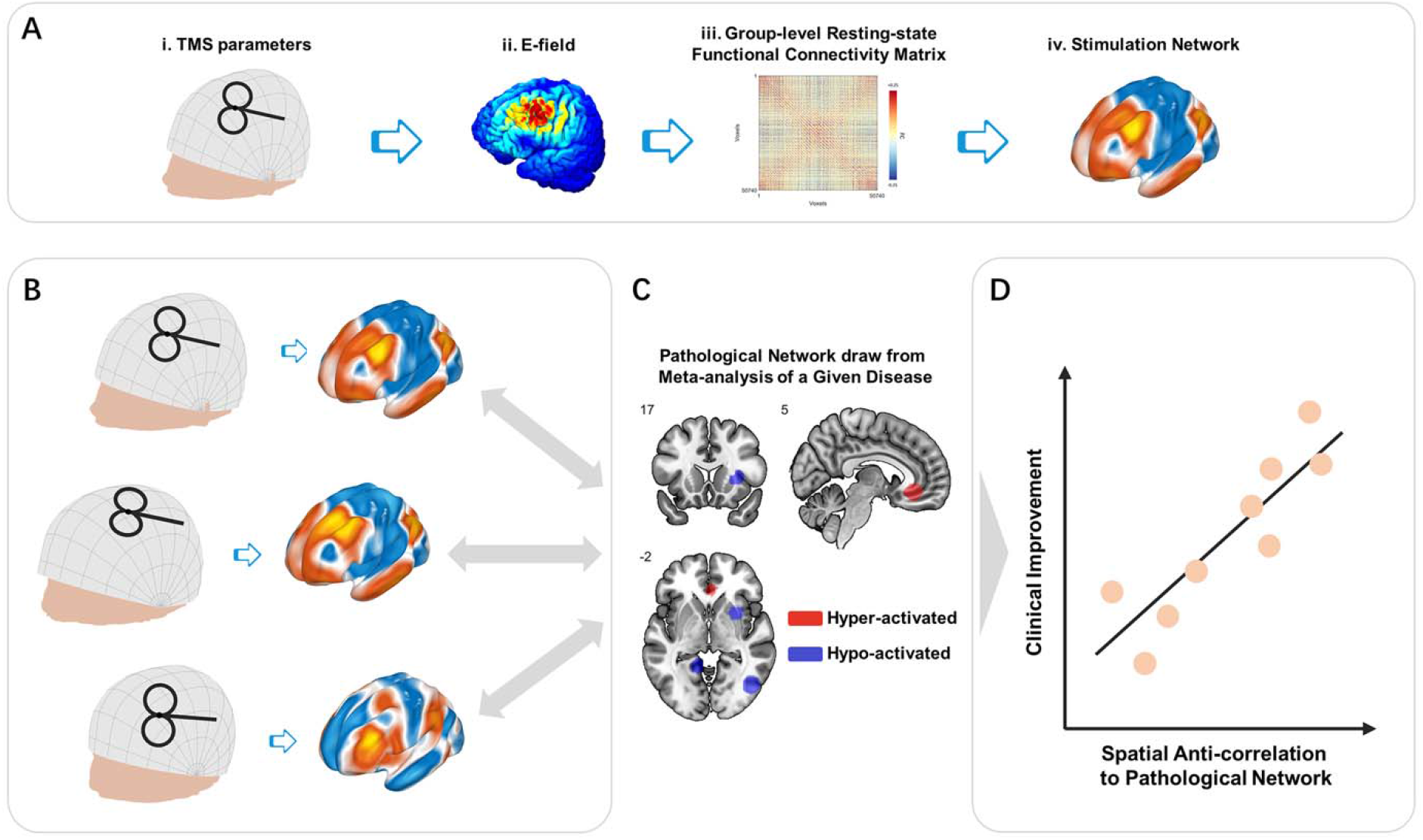
Schematic illustration of network targeting model. **(A)** Stimulation network. For TMS administrated with a given combination of parameters (i), the generated E-filed (ii) defines direct TMS effects on the local cortical region. Group-level rsFC (iii) provides a visualization of the functional network affected via the stimulated cortical region, i.e. the stimulation network (iv). **(B)** Stimulation networks vary among individuals due to both the coil setting and geometry and productivity of individuals’ intra-cranial tissues. **(C)** Comparing to the pathological network of a given disease, **(D)** stimulation networks showing spatial anti-correlation are hypothesized to be associated with better clinical improvement induced by TMS (Fox et al., 2014a).

#### 2.1.2. Parameter Space

We utilized a scalp geometry-based parameter space that describes any possible TMS coil placement with two key parameters (position *s* and orientation θ) on the individual scalp surface (Jiang et al., 2022). The description of position *s* is a pair of coordinates (*p*_NZ_, *p*_AL_) on a continuous proportional coordinate system (CPC), where *p*_NZ_ indicates the position along nasion to inion direction, *p*_AL_ indicates the position along with left preauricular point (AL) to right preauricular point (AR) direction, and (*p*_NZ_ and *p*_AL_) ∈[0 1]×[0 1] (Xiao et al., 2018). The coil orientation (of the handle) is defined in the tangent plane of position *s*. There are two steps to define the direction of orientation 0°. First, we find the intersecting line between the tangent plane and the plane through position *s*, AL, and AR. Second, the 0° direction originates from position *s*, perpendicular to the intersecting line, and points backward. The description of orientation θ is the rotation angle from orientation 0° to the coil handle. For clockwise rotation, θ ∈(−180° to 0°). For anti-clockwise rotation, θ ∈(0° to 180°]. In practice, both parameters of *s* and θ can be implemented with manual measurement (Jiang et al., 2022) and computer-assistant navigation (Xiao et al., 2018; Jiang et al., 2022).

#### 2.1.3. Local effects of TMS stimulation

For a given location and orientation, the local region affected by the TMS induced E-field was estimated by applying FEM modeling on the individual’s T1 image. The FEM modeling was implemented using SimNIBS (Thielscher et al., 2011). According to putative assumptions on the TMS excitatory/inhibitory mechanism, TMS induces an excitatory effect when the pulses are repeatedly delivered at a high frequency (HF) of >5 HZ, while an inhibitory effect is induced at a low frequency (LF) of ≤1 HZ (Pascual-Leone et al., 1998; Dayan et al., 2013).

Such an excitatory/inhibitory effect is limited to the E-field region under coil *para (s*, θ). Assuming a brain with *N* voxels in standard brain space, the local effect of TMS stimulation can be described by an *N*-by-1 vector *E*_*l*_.

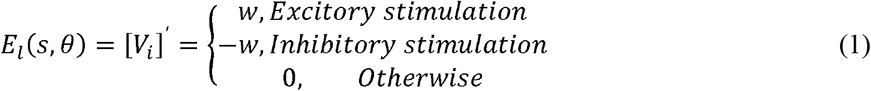

Here, *V*_*i*_ is the local effect of TMS induced on the i^th^ voxel in the E-field region, and *w* is the weight of E-field strength.

#### 2.1.4. RS-FC Profile of TMS Stimulation (Stimulation Network)

In the current model, the rsFC profile of the stimulated region was estimated from the group-level rsFC matrix of the healthy cohort (Weigand et al., 2018). Specifically, the regional rsFC profile, i.e. the ‘stimulation network’, was calculated from the weighted average of whole-brain rsFC seeded from each voxel within the E-field region. The stimulation network corresponding to *para (s*, θ) is given by:

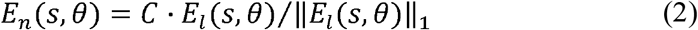

Here, *C* describes the voxel-wise rsFC matrix, and El(s, *θ*) is the local effect of *para (s*, θ), and ∥ ∥_1_ is the 1-norm of a vector, such that E-field weight of suprathreshold voxels sum to one. For *N* gray-matter voxels in MNI space, *C* is given by:

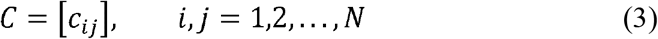

where *c*_*ij*_ is the signed rsFC strength between voxels *i* and *j*.

In the current study, the group-level rsFC was estimated from high-resolution T1 MR images and 8-min resting-state fMRI data of 512 healthy young adults [225 females, age 20.12±1.28 years] from the SLIM database (Liu et al., 2017). The processing of MRI data is detailed in the supplementary materials.

#### 2.1.5. Network Targeting Accuracy

In the proposed model, the metabolic hypo-/hyper-activity was taken as the biological marker for the pathological network of a particular psychiatric disorder. To describe the pathological network, we utilized an image generated from the coordinates-based meta-analysis (CBMA) contrasting a cohort of patients vs. healthy controls (Kühn and Gallinat, 2012; Fox et al., 2014b; Gray et al., 2020). Assuming that the whole gray matter of the brain consists of N voxels in its functional image, which constitute a brain network, the combined activity of these brain voxels represents a state of the brain. The brain states of the patients and controls are represented in N×1 vectors *I*_*pt*_ and *I*_*hc*_, respectively, and the difference between the two states is:

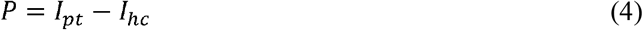

According to the finding that excitatory/inhibitory stimulation on negative/positive FC from the local ROI to deep pathological nodes is beneficial to TMS efficacy (Fox et al., 2014a), we extended this principle by defining the spatial anti-correlation between the pathological network and the TMS stimulation network as the network targeting accuracy (NTA), which we hypothesize can predict the treatment outcome of TMS. For the given para (*s*, θ), the NTA can be quantified by:

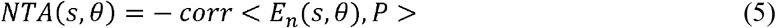

In the current study, we separately utilized the results of two recent CBMA studies as the descriptions of pathological networks for MDD (Gray et al., 2020) and AVH (Kühn and Gallinat, 2012).

### 2.2. Proof-of-concept validation

We conducted three validation experiments to evaluate the feasibility of the proposed NTA model in predicting TMS efficacy from the coil parameters.

First, we tested whether NTA explains the equation-based efficacy (Herbsman et al., 2009; Fox et al., 2012) of empirical DLPFC sites used in treatment of MDD (Rajkowska and Goldman-Rakic, 1995; Paus et al., 2001; Herwig et al., 2003; Okamoto et al., 2004; Cho and Strafella, 2009; Fitzgerald et al., 2009; Herbsman et al., 2009; Rusjan et al., 2010; Fox et al., 2012; Weigand et al., 2018; Cash et al., 2019), we simulated coil setting on T1 images of 68 depression patients [49 females, age 23.69 ± 8.17 years] obtained from OpenNeuro (Anna Manelis et al., 2021; Liuzzi et al., 2021). We calculated site-wise NTA and compared them to the expected efficacy estimated by Herbsman’s equation (Herbsman et al., 2009).

Second, to confirm that the NTA model is capable of predicting the efficacy in the clinical treatment of MDD, we conducted a retrospective validation on a cohort of 33 MDD patients [20 females, age 47.70 ± 7.54 years] who received a two-week treatment of 10 Hz high-frequency rTMS in a previous study (Paillère Martinot et al., 2010). Treatment was targeted using the 5-cm rule or PET-based navigation. We split the 33 patients into two groups (Fox et al., 2012), the left PFC group (*N* = 27) and the right PFC group (*N* = 6). We implemented the NTA model on each patient’s T1 image and calculated NTA from the recorded coil parameters. The calculated NTA was correlated with the actual clinical improvement in each group.

Finally, to test whether the NTA model can be generalized to diseases other than MDD, we conducted another retrospective validation on a cohort of 15 AVH patients [7 females, age 32.07 ± 6.79 years] who received 10 days of 1-Hz rTMS treatment (Paillère-Martinot et al., 2017). Treatment was targeted using fMRI-based navigation. Again, we implemented the NTA model on each patient’s T1 image and calculated NTA from the recorded coil parameters. The calculated NTA was correlated with the actual clinical improvement of each patient.

The full methodology is detailed in the supplementary materials.

### 2.3. Individualized parameter optimization

Motivated by the results of the above analyses, which showed that NTA is able to predict TMS treatment efficacy from the coil parameters, we propose that NTA may serve as an objective function for the individualized optimization of coil parameters. We conducted simulation experiments to demonstrate how optimal parameters vary across patients.

Simulation experiments were conducted on the MDD and AVH cohorts (Paillère Martinot et al., 2010; Paillère-Martinot et al., 2017). In each cohort, we defined a cranial search space covering traditional TMS sites for the two diseases. For MDD the search space had 125 positions × 12 orientations and covered a broad area of left DLPFC (Lefaucheur et al., 2014; Xiao et al., 2018; Cash et al., 2020; Balderston et al., 2021). For AVH, the search space had 122 positions × 12 orientations and covered a broad area including left STG and left TPJ, which have been adopted in TMS treatments for AVH (Hoffman et al., 2003, 2013; Klirova et al., 2013; Lefaucheur et al., 2014; Paillère-Martinot et al., 2017; Xiao et al., 2018). We calculated disease-specific NTA values for each of the parameter combinations, and define the individualized optimal TMS parameters as the combination with maximum NTA.

The full methodology is detailed in the supplementary materials.

## 3. Results

### 3.1. Correlation Between NTA and Equation-Based Clinical Efficacy

To test the hypothesis that NTA predicts treatment efficacy for MDD, we compared NTA and the expected treatment efficacy among 12 TMS sites used for treating MDD (Figure 2A), sourced from previous reviews (Fox et al., 2012; Cash et al., 2020). For each of the cortical targets, the corresponding scalp position was first identified by finding the scalp position with a normal vector pointing to the cortical target, then orientation was fixed at 45° from the mid-line (Fitzgerald et al., 2003; Thomson et al., 2013)(Figures 2B and Figure S1). The parameters of the coil were therefore simulated on each of the 68 individuals from the first cohort. For each cortical site, the across-individual distribution of NTA is shown in Figure 2C, and the mean NTA was used to predict the treatment efficacy estimated with Herbsman’s equation (Herbsman et al., 2009). Across stimulating sites, the NTA showed a significant correlation with HDRS total improvement (*N* = 68, *r* = 0.923, *p* = 9.32×10^−6^, one-tailed) and explained about 85% of the variance assessed by HDRS (Figure 2D). Furthermore, such predictiveness was significantly higher than network targeting models based on randomly generated networks (10^5^ permutation runs, *p* = 0.0343, Figure S2) and was significantly higher than prediction based on randomly reassigned clinical outcomes (10^5^ permutation runs, *p* = 3×10^−5^, Figure S3). Additionally, the estimated NTA is stable when the E-field threshold varied in a range of 75%-99% (r > 0.9, Figure S4) and when the radius of the pathological network foci varied in a range of 4mm-16mm (r > 0.9, Figure S5).

**FIGURE 2.**
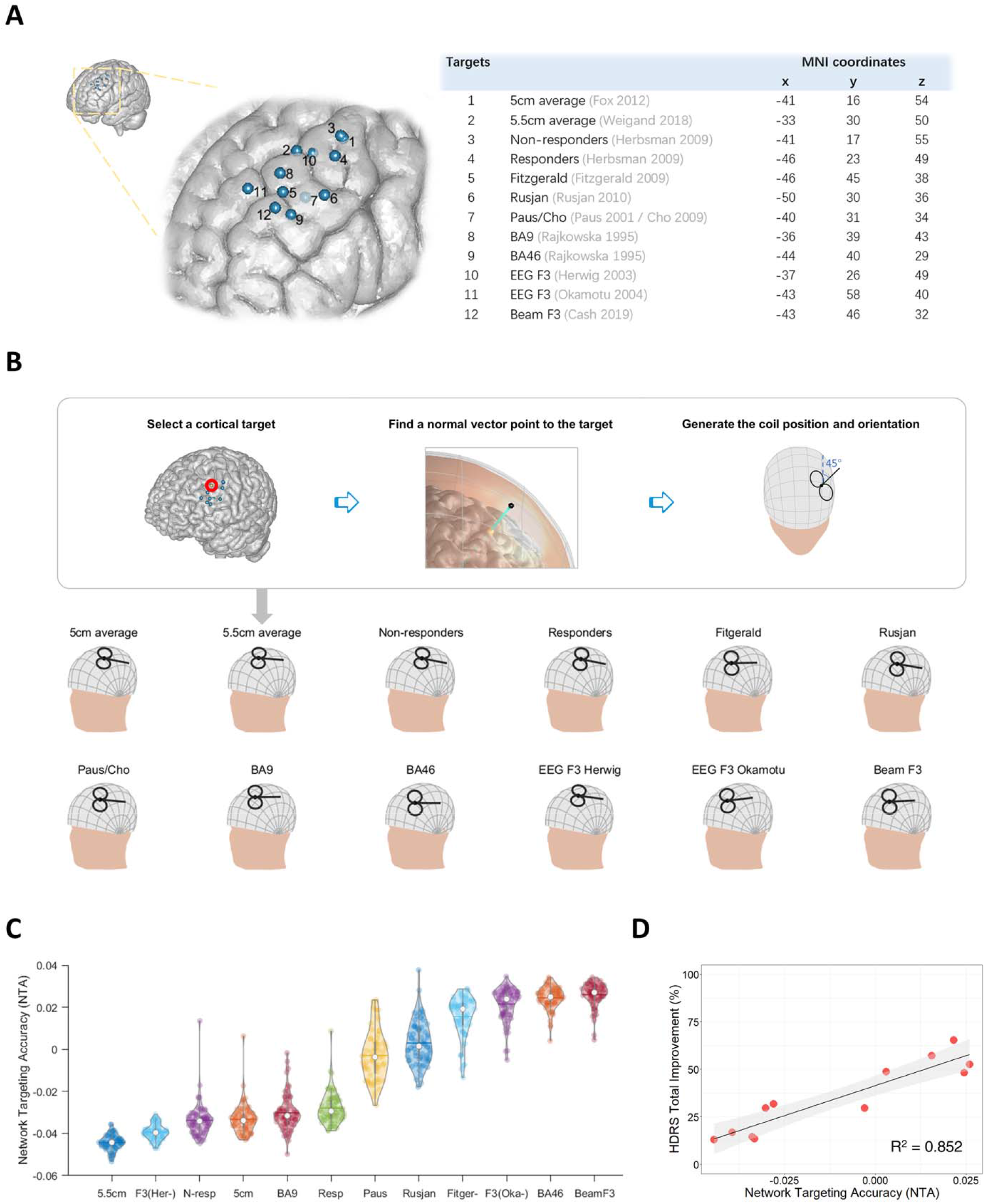
Network targeting model predicts the equation-based TMS treatment efficacy at empirical DLPFC sites in a large depression cohort. **(A)** Empirical target sites of MDD are shown in MNI-152 (Fonov et al., 2011). **(B)** Restoration of TMS parameters from targeted cortical sites. **(C)** NTA of empirical sites across different individuals, each represented with a colored dot (*N* = 68). **(D)** Correlation between the average NTA and the equation-based HDRS total improvement (*p* = 9.32×10^−6^, one-tailed).

### 3.2. Correlation Between NTA and Treatment Efficacy on MDD Patients

In the MDD cohort who received TMS treatment, the recorded TMS coil positions and orientations are shown in Figure 3A. Across stimulating sites in left PFC (Fox et al., 2012), NTA showed a significant correlation with MADRS total improvement (*N* = 27, *r* = 0.337, *p* = 0.043, one-tailed) and explained about 11% of the variance assessed by MADRS (Figure 3B). Furthermore, such predictiveness was significantly higher than network targeting models based on randomly generated networks (10^5^ permutation runs, *p* = 0.0306, Figure S2) and was significantly higher than prediction based on randomly reassigned clinical outcomes (10^5^ permutation runs, *p* = 0.0355, Figure S3).

**FIGURE 3.**
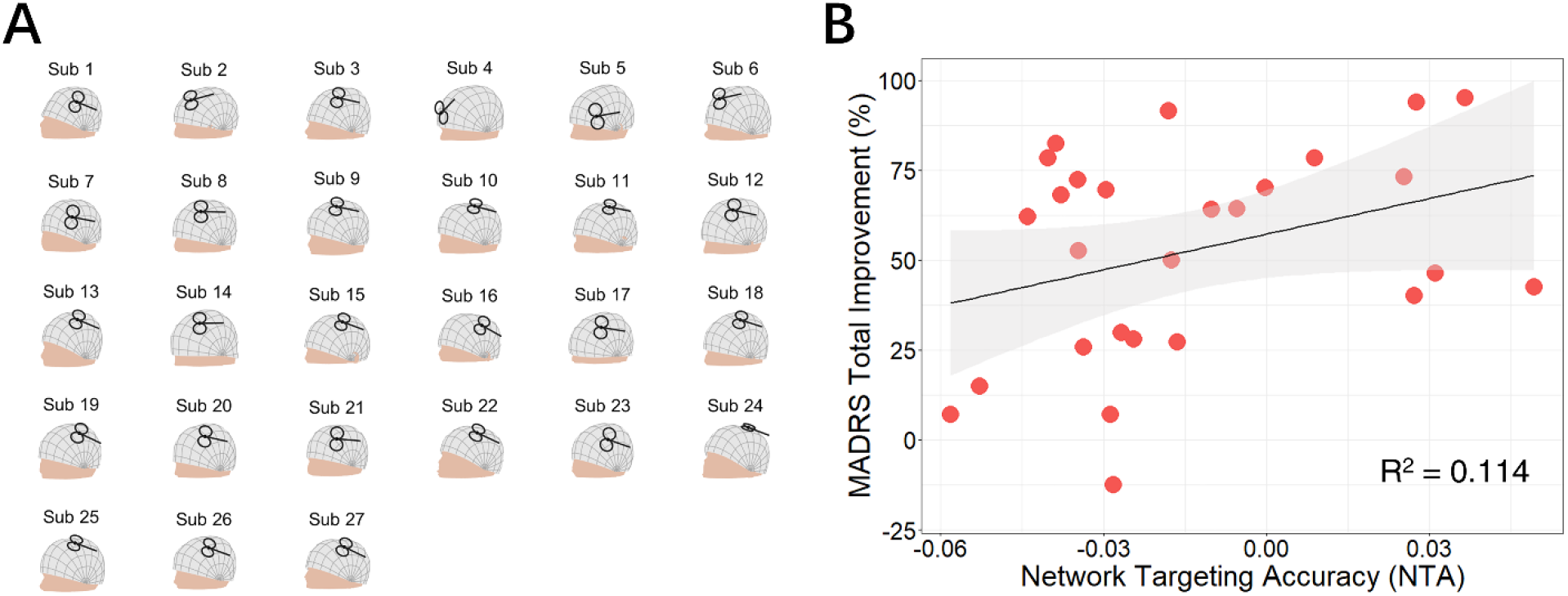
Network targeting accuracy predicts treatment efficacy in the clinical MDD cohort. **(A)** Coil placement of left PFC patients shown on individual head models. **(B)** Correlation between NTA and MADRS total improvement (*N* = 27, *p* = 0.043, one-tailed).

Additionally, NTA was stable when the E-field threshold varied in a range of 75%-99% (r > 0.9, Figure S4) and when the radius of the pathological network foci varied in a range of 4 mm-16 mm (r > 0.9, Figure S5).

The predictiveness of the NTA model was limited within the left PFC. For the six other patients in this cohort who received high-frequency TMS treatment in the right PFC, their clinical outcome was not predicted by the NTA model (*N* = 6, *r* = −0.310, *p* = 0.725, one-tailed, Figure S6). This result may due to that the treatment outcome in these subjects come from a placebo effect rather than the TMS modulation, given evidence that the anti-MDD efficacy of high-frequency rTMS is specific to left DLPFC (Lefaucheur et al., 2014).

### 3.3. Correlation Between NTA and Treatment Efficacy on AVH Patients

TMS coil positions and orientations of the active group are shown in Figure 4A. Across stimulating sites, NTA showed a significant correlation with AHRS total improvement (*N* = 15, *r* = 0.556, *p* = 0.016, one-tailed) and explained about 31% of the variance assessed by AHRS (Figure 4B). Furthermore, such predictiveness was significantly higher than network targeting models based on randomly generated networks (10^5^ permutation runs, *p* = 0.0042, Figure S2) and was significantly higher than prediction based on randomly reassigned clinical outcomes (10^5^ permutation runs, *p* = 0.0176, Figure S3). Additionally, the estimated NTA was stable when the E-field threshold varied in a range of 75%-99% (r > 0.8, Figure S4) and when the radius of pathological network foci varied in a range of 4mm-16mm (r > 0.9, Figure S5).

**FIGURE 4.**
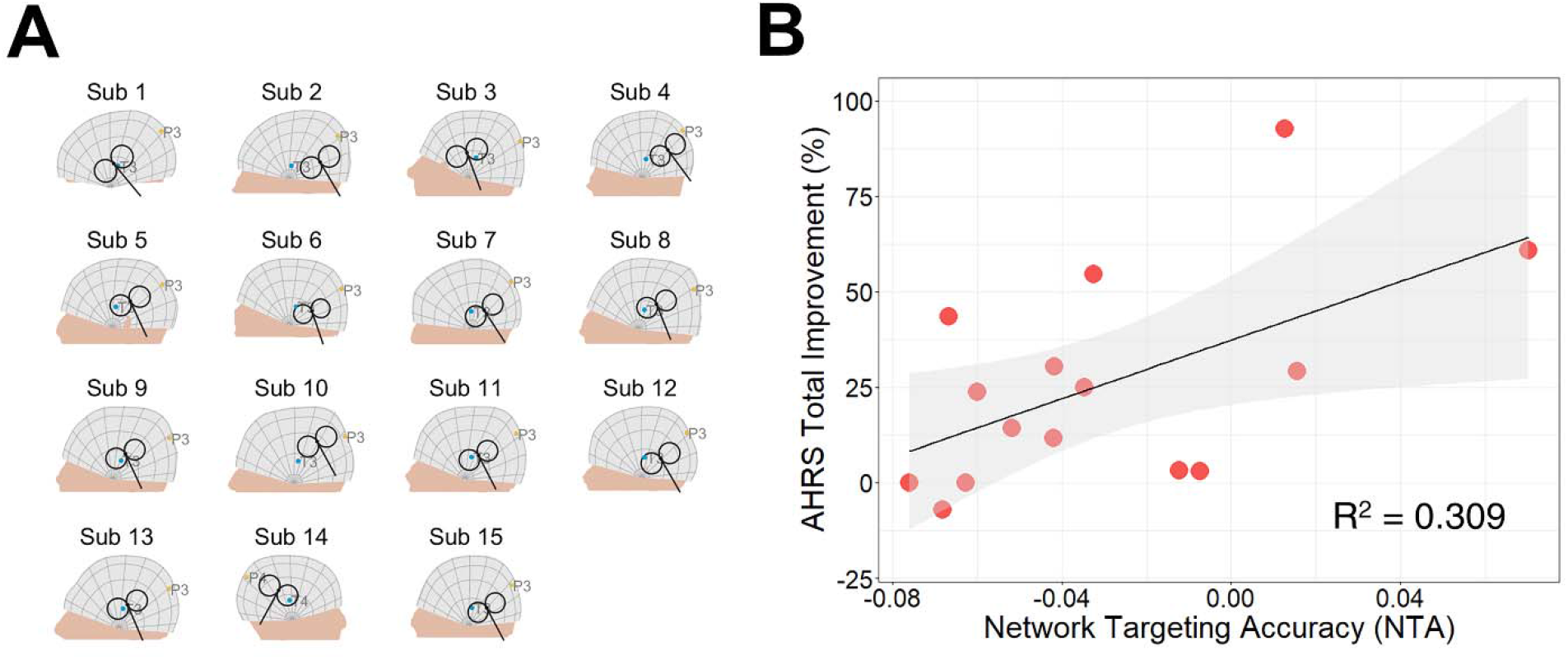
Network targeting accuracy predicts treatment efficacy in the clinical AVH cohort. **(A)** Coil placement of active group patients shown on individual head models. **(B)** Correlation between NTA and AHRS total improvement (*N* = 15, *p* = 0.016, one-tailed).

We further correlated NTA with changes in other clinical assessments, including scales of schizophrenia (SAPS and SANS)(Table 1). First, the predictiveness of NTA showed specificity to TMS induced changes in positive symptoms (*N* = 15, *r* = 0.572, *p* = 0.013, one-tailed) but not in negative symptoms (*N* = 15, *r* = 0.021, *p* = 0.470, one-tailed). Second, within the sub-scales of SAPS, NTA predicted changes in hallucination-related items, but not in other items related to delusion, bizarre behavior, and positive formal thought disorder. Collectively, the above results indicate predictiveness of NTA is specific to the targeted symptom.

**TABLE 1.**
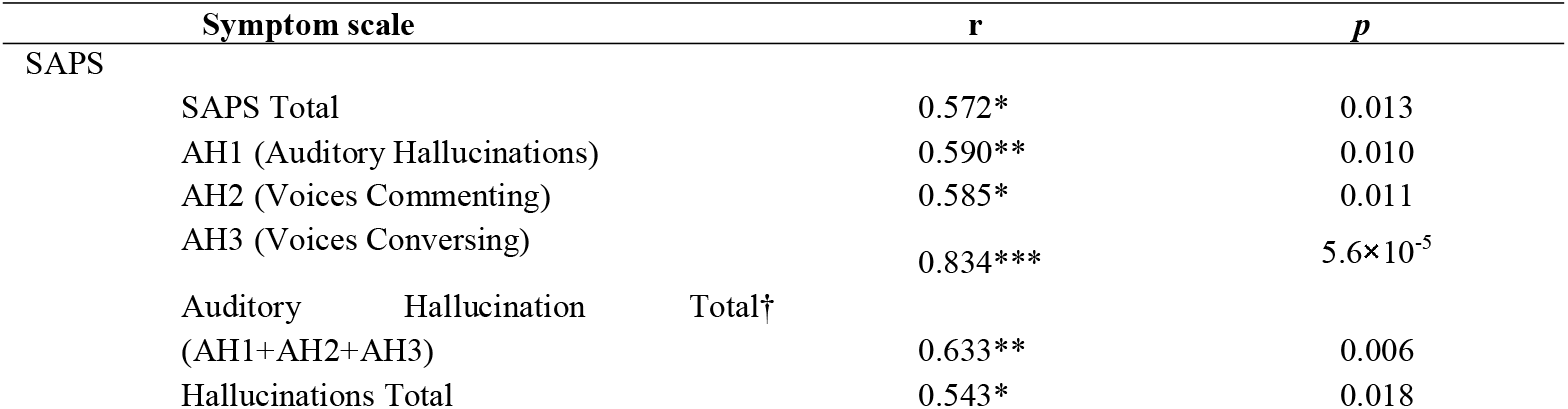
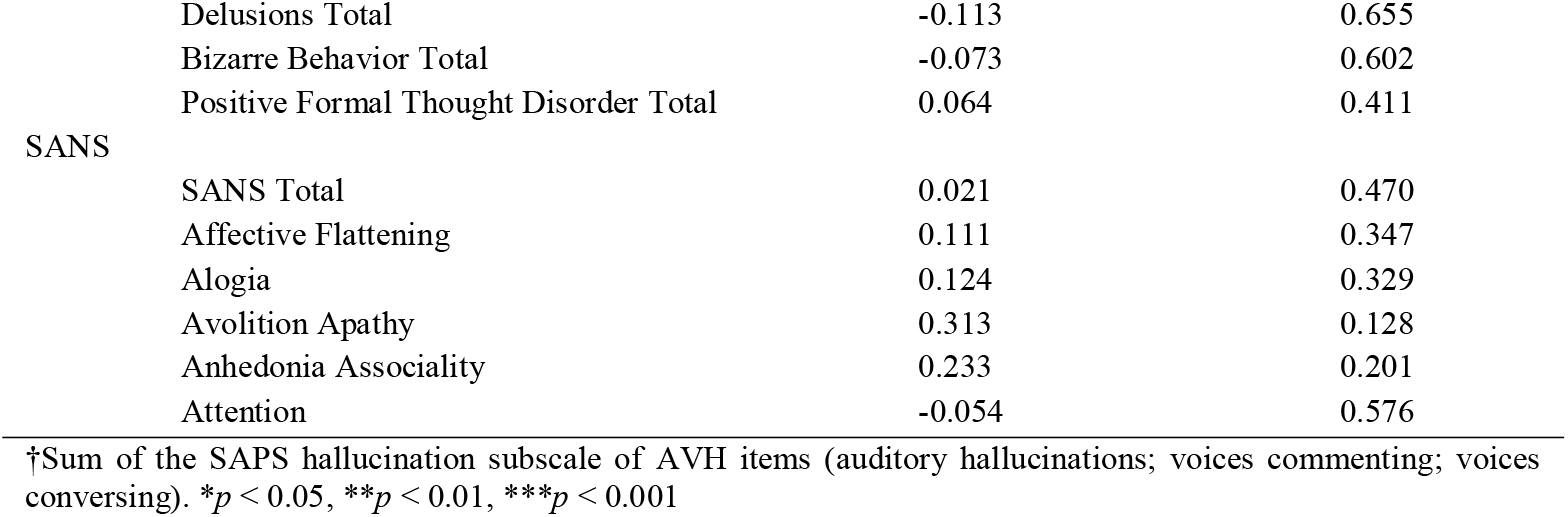
Symptom specificity of predictions from the NTA model.

### 3.4. Position-orientation Interaction on Estimated Treatment Efficacy and Individualized Optimization

In the MDD cohort, we simulated the NTA model for MDD on each patient within the left DLPFC (Figure 5A). Possible combinations of position and orientation formed a 2-D parameter space which was subdivided into a 125-by-12 (position by orientation) grid. We calculated the estimated NTA for each of the combinations. Across the 27 individuals, both the position (*F*(124, 38974) = 375.490, *p* < 0.001) and orientation (*F*(11, 38974) = 4.201, *p* < 0.001) had significant main effect on NTA; there was also a significant interaction effect (*F*(1364, 38974) = 16.766, *p* < 0.001) between the two parameters. Within the left DLPFC, the optimal parameter was defined as the combination with the highest value of NTA (Figure 5B). Optimal parameters varied across different individuals (Figures 5C,D).

**FIGURE 5.**
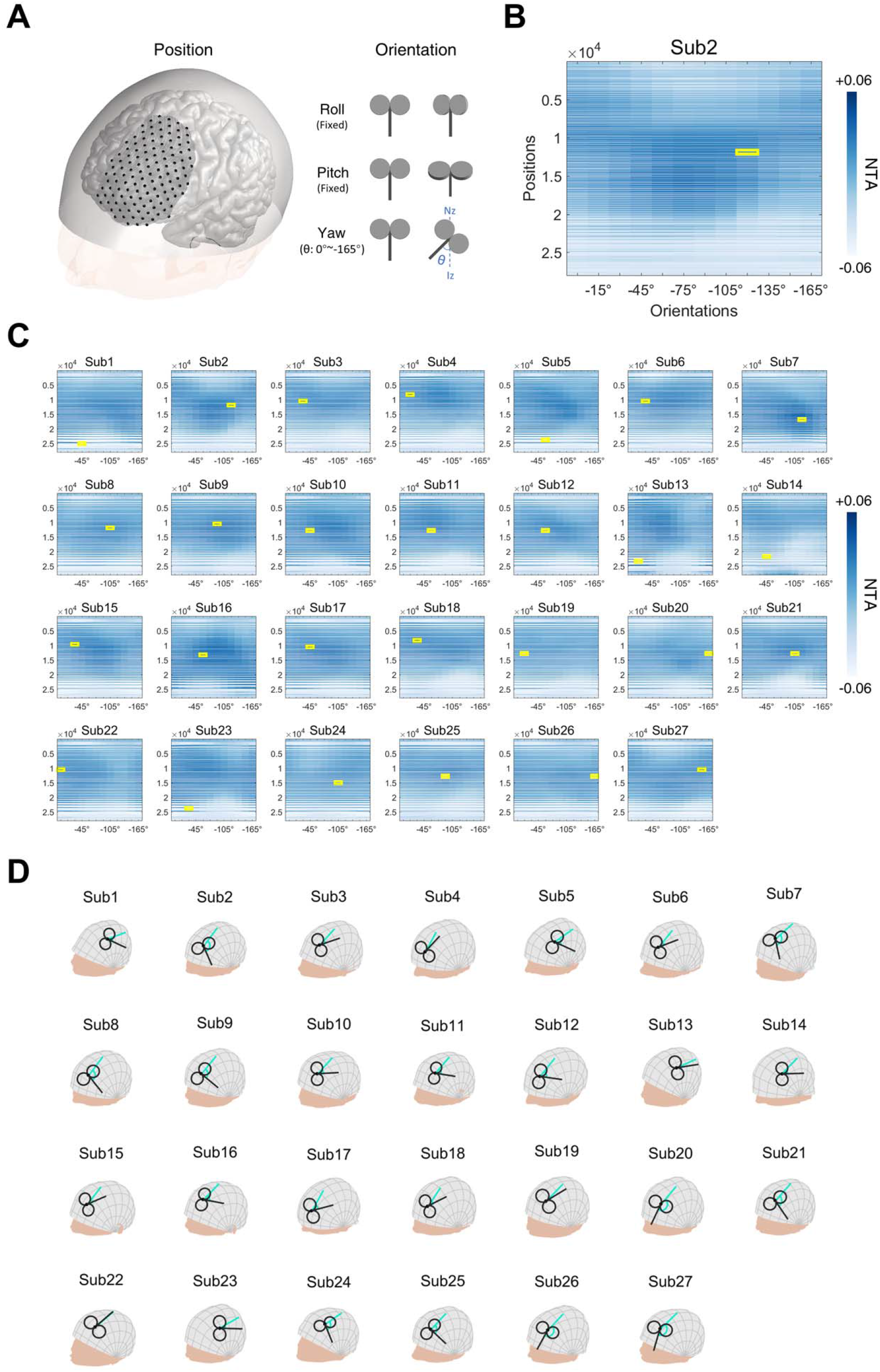
MDD Simulation Experiment. **(A)** Illustration of positions and orientations of a representative individual. Large black dots represent 125 positions in the search space. For each position, 12 coil orientations, in the normal plane at the position (0° ∼ −165°, 15-degree intervals), were tested. NTA was calculated for each pair of position and orientation. **(B)** NTA value distribution in the search grid. Each position in the 2-D grid represents a combination of position and orientation. **(C)** Maximum NTA was found in all patients (yellow border). Search space was interpolated from 125×12 to 27977 × 12 for visualization purposes. **(D)** The optimal TMS coil placements are shown in individual scalp spaces. The Cyan arrow represents 0° at each position.

In the AVH cohort, we performed a similar simulation on a 122-by-12 (position by orientation) parameter space covering left STG and left TPJ, places where TMS is commonly administrated (Figure 6A). Again, we found significant main effects in both parameters of position (*F*(121, 20482) = 102.572, *p* < 0.001) and orientation (*F*(11, 20482) = 11.146, *p* < 0.001), and interaction between the two parameters (*F*(1331, 20482) = 9.220, *p* < 0.001). Figure 6B illustrates the distribution of NTA and optimal parameters in a representative individual. Optimal parameters also varied among different individuals (Figures 6C,D).

**FIGURE 6.**
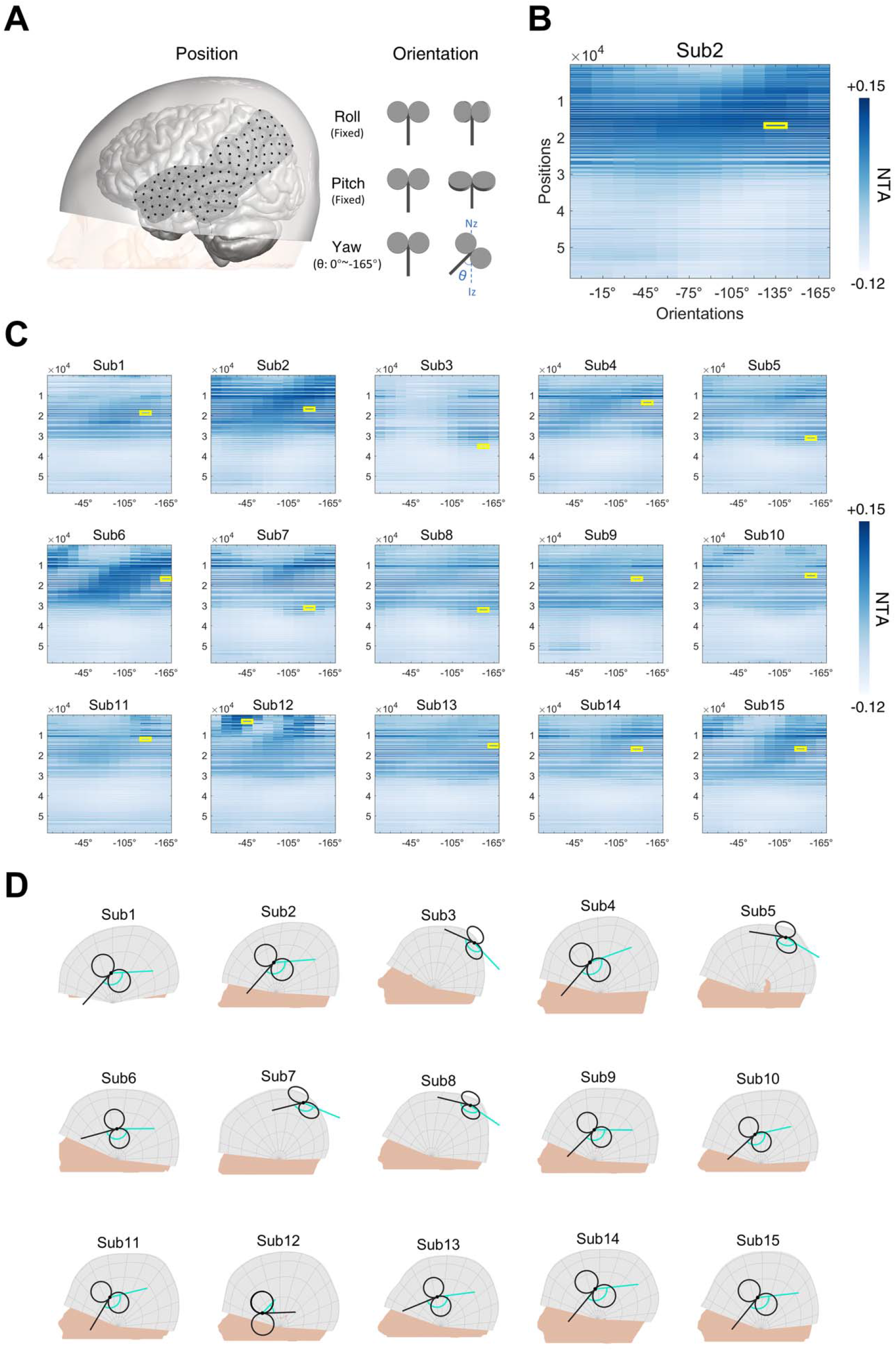
AVH Simulation Experiment. **(A)** Illustration of positions and orientations of a representative individual. Large black dots represent the 122 positions in the search space. For each position, 12 coil orientations (0° ∼ −165°, 15-degree intervals) were tested. NTA was calculated for each pair of position and orientation. **(B)** NTA value distribution in the search grid. Each position in the 2-D grid represents a combination of position and orientation. **(C)** maximum NTA found in all patients (yellow border). Search space was interpolated from 122×12 to 58470 × 12 for visualization purposes. **(D)** The optimal TMS coil placements are shown in individual scalp spaces. The Cyan arrow represents 0° at each position.

## 4. Discussion

In this work, we proposed a novel network targeting model for guiding individualized TMS coil settings for the treatment of psychiatric disorders. The model linked the TMS parameter space of coil position and orientation with the improvement of clinical symptoms after treatment, with a hypothesis that the treatment outcome was associated with the extent of modulation by TMS on the whole pathological network of a given disease. For a proof-of-concept, the proposed model was validated by retrospectively predicting the expected efficacy at empirical DLPFC sites based on a large depression cohort and the outcome of two clinical cohorts (MDD and AVH) that received TMS treatments. The proposed model significantly predicted treatment efficacy from the position and orientation of TMS parameters. Furthermore, in the AVH cohort, the prediction was both specific to the symptom corresponding to the targeted pathological network. Finally, we further applied the model to individual optimization of TMS parameters within the search space of traditional MDD and AVH treatment on the scalp. The results of optimization showed the variance of optimal individual parameters and the interaction of position and orientation.

Consistent with related previous studies, our results demonstrated that considering both the local ROI and the related functional circuit affected by rTMS is a potential way to inform an accurate modulation for psychiatric disorders, in comparison to the traditional ROI-based approach. In a series of seminal studies in MDD, research has shown that the stimulation ROI of DLPFC with stronger anti-correlation with SGC tends to show better clinical improvement (Fox et al., 2012; Weigand et al., 2018; Cash et al., 2021a). While the mechanism is still unknown (Mayberg, 1997; Speer et al., 2000; Li et al., 2004; Padberg and George, 2009; Kito et al., 2011; Fox et al., 2012, 2014a; Philip et al., 2018), the fact that SGC and DLPFC are two critical regions belonging to the frontal-limbic network, the putative pathological network of MDD identified by various neuroimaging studies, suggests that the information about the whole pathological network is necessary to inform effective TMS treatment. In line with this notion, the proposed model generalized the single SGC-DLPFC connectivity method to the collective effect of the whole pathological network. In addition, we compared the SGC-DLPFC method with the whole pathological network targeting method. The predictiveness was improved in the method targeting the whole pathological network although to a limited extent (Figure S8), suggesting that the SGC-DLPFC circuit may still play a dominant role in guiding TMS treatment for MDD. Compared to the hypothesis-driven method based on a specific ROI such as SGC for MDD, the data-driven network targeting model is particular valuable for generalizing the prediction of treatment outcomes from MDD to other psychiatric disorders such as AVH.

For TMS-based treatment of AVH, traditional targeting strategies are mainly based on a single-ROI target within the left temporoparietal cortex, either defined by anatomical landmarks such as TP3 (Hoffman et al., 2003) or left Wernicke (Hoffman et al., 2013), or functional foci showing abnormal activation (Sommer et al., 2007). Though techniques like neuronavigation have increased the accuracy in locating these ROIs, improvement in treatment efficacy is relatively limited (Slotema et al., 2011). Regarding this point, our retrospective analysis showed that minimizing the spatial distance to the targeted ROI was not related to treatment efficacy (Figure S9). Instead, minimizing the functional distance to the pathological network of AVH was shown to be a potential goal for optimization.

The interaction of position and orientation suggests the necessity of individual optimization. In the simple case, the MEP is highly dependent on coil position and orientation and an individual’s intracranial anatomy (Windhoff et al., 2013; Laakso et al., 2014; Reijonen et al., 2020). In a more complicated case, the combination of coil position and orientation affects the targeting of functional networks (Opitz et al., 2016). In line with these studies, the proposed network targeting model also showed a significant interaction between coil location and orientation on NTA. This suggests the necessity of including the coil orientation in both the parameter space and the individualized optimization process based on individual structural images.

In estimating the stimulation network of TMS, our results support the utility of group-level functional connectomes, as suggested in previous studies of similar functional connectome-based approaches (Fox et al., 2012, 2014a; Weigand et al., 2018; Cash et al., 2019). It is worth noting that other evidence also suggests that the treatment efficacy of rTMS may be further improved by customizing stimulation sites based on individual differences in functional connectivity (Fox et al., 2013; Cash et al., 2019, 2021b). However, compared with individual functional connectivity, the advantage of using the normative connectome data is the generally higher signal-to-noise ratio. Data acquired on the normative population can be optimized by using improved technologies of acquisition, enlarging the sample size (Van Essen et al., 2012), and increasing the density of sampling in individuals (Laumann et al., 2015), which are usually difficult to conduct on patient populations (Horn and Fox, 2020). The trade-off between meaningful individual differences and the quality of functional connectivity data remains to be addressed in future work.

The proposed model derives the pathological network from the contrast of patient vs. healthy control. An implication is that reducing the biological deviation of the patient cohort from the healthy is a feasible direction for optimizing the parameters of TMS when treating mental disorders. Within such a model, further improvement can be made in several directions. This study used the altered baseline metabolic pattern of patients relative to healthy controls as the neural target for TMS-based treatment. As promising alternatives, symptom-specific pathological networks, compensatory networks, and side-effect networks for psychiatric diseases are worth considering in future studies. Psychiatric disorders are often diagnosed by heterogeneous symptoms, of which the biological markers are elusive (Abi-Dargham and Horga, 2016). Current efforts searching for neural markers of psychiatric disorders have identified distinct networks underlying the severity or the response to the treatment of psychiatric symptoms (Drysdale et al., 2017; Siddiqi et al., 2020). Therefore, nodes of these networks would be potential targets for the development of symptom-specific treatments. An interesting line of research focuses on identifying networks associated with treatment-induced side effects (Horn and Fox, 2020), and the results might be integrated into the proposed model as a “to-avoid” network in planning treatment. Apart from searching nodes of the pathological network, Balderston used a data-driven approach to link rsFC and symptoms of depression (Balderston et al., 2021), demonstrating the feasibility of edge-based targeting in TMS treatment. Such an edge-based pathological network will be considered in our model in the future.

There are several limitations to the current work. First, the sample size for the validation experiment was small. Therefore, the correlation analysis based on such a small sample might be unstable and provide a biased estimation of the true effect size. Second, the retrospective validation might be confounded by factors insufficiently controlled, e.g., variance in TMS protocols or heterogeneity of patients. Therefore, prospective validation would be necessary for follow-up research. Third, though the proposed NTA model showed its ability to generalize to AVH, a disease other than MDD, from which the core idea of the model arose, whether it can generalize to other psychiatric diseases need to be further investigated. Fourth, the current NTA model focused on TMS coil position and orientation, which are a subset of the TMS parameters. Other dimensions of the full parameter space such as the number of pulses, stimulation intensity, and temporal patterns of the pulses (Lefaucheur et al., 2014) need to be considered in future studies.

## 5. Conclusion

This study proposed a novel network targeting model for guiding individualized TMS treatment of psychiatric disorders. For a proof-of-concept, retrospective validation on MDD showed that the proposed model was capable of predicting clinical outcomes from TMS placement settings. The model showed comparable predictiveness for AVH, demonstrating its generalizability. Finally, the proposed model showed potential for guiding individualized TMS placement. Though prospective validation is needed, this network targeting model may offer an opportunity for improving the current TMS-based treatment of psychiatric disorders.

## Conflict of Interest

The authors declare that the research was conducted in the absence of any commercial or financial relationships that could be construed as a potential conflict of interest.

## Author Contributions

**Zhengcao Cao**: Conceptualization, Formal analysis, Methodology, Investigation, Visualization, Data Curation, Software, Writing - original draft. **Xiang Xiao**: Conceptualization, Formal analysis, Investigation, Methodology, Software, Writing - original draft. **Yang Zhao**: Formal analysis, Methodology. **Yihan Jiang**: Writing -review & editing. **Cong Xie**: Software. **Marie-Laure Paillère-Martinot**: Resources, Writing - review & editing. **Eric Artiges**: Resources. **Zheng Li**: Writing - review & editing. **Zafiris J. Daskalakis**: Writing - review & editing. **Yihong Yang**: Funding acquisition, Conceptualization, Supervision, Writing - review & editing. **Chaozhe Zhu**: Funding acquisition, Conceptualization, Supervision, Writing - review & editing, Conceptualization.

## Funding

This work was supported by the National Natural Science Foundation of China (grant no. 82071999). X.X. and Y.Y. are supported by the Intramural Research Program of the National Institute on Drug Abuse, the National Institute of Health, USA.

## Acknowledgments

INSERM is acknowledged for sponsorship of the AVH cohort acquisition. INSERM U A10 holds the copyright of the AVH cohort. Jean - Luc Martinot, MD, Ph.D., INSERM U A10, is acknowledged for setting up the sponsorship and contributing to data acquisition in this patient cohort.

## Data Availability Statement

The cohorts, including the structure and resting-state functional MRI, used to construct the voxel-wise connectome are from the Southwest University Longitudinal Imaging Multimodal (SLIM) database (http://fcon_1000.projects.nitrc.org/indi/retro/southwestuni_qiu_index.html) and is openly available. The list of analyzed participants can be obtained upon request from C.Z. The results of coordinate-based meta-analysis have been reported in studies published previously. The T1 images of patient cohorts for MDD and AVH are not publicly available due to the confidentiality policy of INSERM U A10, but are available upon reasonable request by contacting M.L.P.M.

The code used in the current study for developing the model is available upon reasonable request by contacting C.Z.

## Notes

### Competing Interest Statement

The authors have declared no competing interest.

