## supplementary materials for "Targeting the pathological network: feasibility of network-based optimization of transcranial magnetic stimulation coil placement for treatment of psychiatric disorders"

### Supplementary methods

### 1. Proof-of-concept validation

In summary, one healthy MRI cohort was used to construct the network targeting model and two meta-analysis studies were utilized as coordinate-system based descriptions of the pathological network for MDD and AVH, respectively, in the network targeting model.

Three clinical cohorts were used to validate the proposed network targeting model in separate experiments. In the cohort of depression, we tested the correlation between NTA and equation-based clinical efficacy. In the second cohort of MDD and the third cohort of AVH (patients who received TMS treatment), NTA was calculated from the recorded coil settings and retrospectively compared with the treatment outcome.

#### 1.1. Participants and data acquisition

##### *1.1.1. Healthy cohort for constructing group-level functional connectivity*

The MRI cohort included anatomical MRI and rsfMRI data from 512 healthy adults [225 females, age 20.12 ± 1.28 years]. This was used to generate a group-level rsFC map. The cohort was from the Southwest University Longitudinal Imaging Multimodal (SLIM) database (Liu et al., 2017) of the International Neuroimaging Data-sharing Initiative (INDI). This study was approved by the Research Ethics Committee of the Brain Imaging Center of Southwest University. Informed written consent was obtained from each subject. The high-resolution 3D T1-weighted structural images were obtained using a Magnetization Prepared Rapid Acquisition Gradient-echo (MPRAGE) sequence (TR/TE=1900 ms/2.52 ms, FA=9°, FOV = 256 × 256 $mm^2$; slices = 176; thickness = 1.0 mm; voxel size = 1×1×1 $mm^3$). Eight minutes of resting-state functional data were acquired for each participant. During scanning, participants were asked to close their eyes and rest without thinking about anything but refrain from falling asleep. The functional images were acquired using a gradient echo-planar imaging (EPI) sequence with the following parameters: slices = 32, repetition time (TR)/echo time (TE) = 2000/30 ms, flip angle = 90°, field of view (FOV) = 220 × 220 $mm^2$, and thickness/slice gap = 3/1 mm, and voxel size = 3.4 × 3.4 × 3 $mm^3$. The SLIM study was conducted in accordance with the Declaration of Helsinki revised in 1989.

##### *1.1.2. Clinical cohorts for retrospective validation*

In the first validation experiment, the empirical DLPFC sites and coil orientation were simulated on T1 images of a large depression cohort adopted from the OpenNeuro database (https://openneuro.org). This cohort included a total of 68 depression patients [49 females, age 23.69 ± 8.17 years] from two studies. The first study included 26 major depressive disorder patients and 13 persistent depression disorder patients (Anna Manelis et al., 2021). The second study included 29 major depressive disorder patients (Liuzzi et al., 2021). For the first study (Anna Manelis et al., 2021), T1 images (MPRAGE sequence, voxel size = 1 × 1 × 1 $mm^3$) were acquired on a 3T GT scanner (GE, Milwaukee, USA). For the second study (Liuzzi et al., 2021), T1 images (MPRAGE sequence, 208 contiguous slices, voxel size = 0.8 × 0.8 × 0.8 $mm^3$, FOV = 256, TE = 2.22 ms, flip angle = 8°) were acquired on a 3T Prisma scanner (Siemens, Washington, D.C., USA).

In the second validation experiment, we utilized data from a previous study (Paillère Martinot et al., 2010). One patient was excluded from the analysis due to the failure of segmentation of the T1 image. The clinical MDD cohort in the current analysis included 33 patients [20 females, age 47.70 ± 7.54 years]. These patients with a DSM-IV-R diagnosis of major depressive disorder were recruited by senior psychiatrists from consecutive admissions at five university psychiatry departments [Montgomery–Asberg Depression Rating Scale (MADRS) 32.85 ± 7.51]. Exclusion criteria included age >65 years, alcohol or substance dependence in the past 6 months, electroconvulsive therapy (ECT) treatment in the past 6 months, any present medical condition, history of epileptic seizures, history of neurological disorders or substantial brain damage, and contraindication to magnetic fields, according to established safety criteria (Eric M. Wassermann, 1998). These patients underwent a two-week treatment of 10 Hz high-frequency rTMS, and 10 sessions of 10-Hz TMS (1600 pulses/session) at a 90% motor threshold. T1 images (3D sequence, 124 contiguous slices, voxel size = $0.94 \times 0.94 \times 1.3$ $mm^3$) were acquired on a 1.5T Signa scanner (GE, Milwaukee, USA). The rTMS was administrated at the most hypometabolic prefrontal area of the patient revealed by PET imaging or was administrated at a site determined by the standard 5-cm rule. For PET imaging, the peak voxel was projected on the scalp using Anatomist software (http://brainvisa.info). Then, the TMS coil was positioned to the scalp target and tightly maintained on the head with a tourniquet (Andoh et al., 2009). For the standard 5-cm rule, the TMS coil was positioned 5 cm anterior to the motor hot spot of the hand. The coil handle was rotated 45° from the midline, and the handle was always backward. The MNI coordinates of individual stimulation locations were recorded in the cohort, and the treatment outcome was assessed with the MADRS. The original study was performed in accordance with the Declaration of Helsinki. The study was approved by the ethics committee Ile de France 6, Paris. Written informed consent was obtained from all subjects after the full description of the study. We split the 33 patients into two group, left PFC group ($N = 27$) and right PFC group ($N = 6$). The left PFC group is chosen in a former study (Fox et al., 2012).

In the third validation experiment, we utilized data from a previous study (Paillère-Martinot et al., 2017). The clinical AVH cohort in the current analysis included 15 AVH patients [7 females, age 32.07 ± 6.79 years]. These patients who met the criteria for a diagnosis of schizophrenia according to a checklist of DSM-IV-TR criteria or schizoaffective disorder were consecutively recruited by senior psychiatrists from three out-patient clinics at university psychiatry departments [Auditory Hallucination Rating Scale (AHRS) 28.67 ± 5.38]. The patients were screened for daily AVH for at least 3 months as assessed by interviews of the patients and their psychiatrists. Exclusion criteria included age over 65, pregnancy, alcohol or substance abuse or dependence in the past 6 months, electroconvulsive therapy in the past 6 months, any present medical condition, history of other psychiatric or neurological disorders, and contraindication to magnetic fields. In the current AVH cohort, 15 patients in the active group were administrated with MRI-guided 1-HZ rTMS for ten consecutive working days. T1 images (3D sequence, 124 contiguous slices, voxel size = $0.94 \times 0.94 \times 1.3$ $mm^3$) were acquired on a 1.5T Signa scanner (GE, Milwaukee, USA). The AVH patients performed a language fragment-detection task. Individual stimulation locations were determined based on the most significant peak voxel in the posterior part of the superior temporal gyrus or, if absent, within the inferior parietal gyrus or the inferior parietal, the middle temporal gyrus. The peak voxel was projected on the scalp using Anatomist software (http://brainvisa.info). Then, the TMS coil was positioned on the scalp target and tightly maintained on the head with a tourniquet (Andoh et al., 2009). The coil handle was pointed downwards, perpendicular to the temporal (left-side: T3 or right-side: T4) and parietal (left-side: P3 or right-side: P4) of EEG points. The MNI coordinates of individual stimulation locations were recorded in the cohort, and the clinical outcomes were measured with the AHRS. Another two clinical scales, the Scale for the Assessment of Positive Symptoms (SAPS) and the Scale for the Assessment of Negative Symptoms (SANS), were also included for analysis. The original study was performed following the Declaration of Helsinki and approved by the local ethics committee Ile-de-France 6, Paris (approval no. 111-01). Written informed consent was obtained from all participants after they completed the description of the study.

#### 1.2. **Calculating network targeting accuracy**

##### ***1.2.1. TMS coil position and orientation***

In the first validation experiment, on each T1 image of depression 68 patients from the OpenNeuro database, we simulated the positioning TMS coil according to MNI coordinates of empirical DLPFC targets (Rajkowska and Goldman-Rakic, 1995; Paus et al., 2001; Herwig et al., 2003; Okamoto et al., 2004; Cho and Strafella, 2009; Fitzgerald et al., 2009; Herbsman et al., 2009; Rusjan et al., 2010; Fox et al., 2012; Weigand et al., 2018; Cash et al., 2019) by setting the coil orientation 45° from the mid-line commonly used in MDD treatment (Fitzgerald et al., 2003; Thomson et al., 2013). We first segmented T1 images of 68 patients with SimNIBS 3.2 (Thielscher et al., 2015;

Saturnino et al., 2019). Then we utilized the head surface nodes of these patients to generate their individual parameter space (Jiang et al., 2022). The MNI coordinates of left DLPFC sites were converted to subject space with the mni2subject_coords function of the SimNIBS 3.2 package. In the individual space, parameters of TMS coil placement were restored as follows: 1. For a given cortical site, we selected CPC on the scalp within 30 mm from the cortical site as candidate scalp positions. 2. From each of the scalp positions, we calculated normal vectors of the scalp surface, and accordingly the distance between the cortical site and the normal vectors. 3. CPC position with minimal distance from the corresponding normal vector to the cortical site was chosen as the simulated coil position. 4. The handle direction was rotated 45° from the midline, and the handle was always pointed backward (Figure 2 & Figure S1). In addition, the coil placements of the cohort are shown in parameter space (Supplementary Table 3 & Supplementary Table 4).

In the second validation experiment, we first segmented T1 images of 33 MDD patients with SimNIBS 3.2. Then we utilized the head surface nodes of these patients to generate their individual parameter space (Jiang et al., 2022). We utilized the projection method in Anatomist software (Andoh et al., 2009). Specifically, the MNI coordinates of individual stimulation locations were converted to subject space with the mni2subject_coords function of the SimNIBS 3.2 package and then were projected on the scalp by calculating the center of gravity of the (n = 10) closest head surface nodes. Coil orientation was quantified according to the recorded stimulation parameter

(Paillère Martinot et al., 2010). We first defined a normal vector that was averaged over a round area around the target site (5 mm of radius). The handle direction was rotated 45° from the midline, and the handle was always pointed backward (Figure 3A). In addition, the coil placements of the cohort are also shown in parameter space (Supplementary Table 5).

In the third validation experiment, we first segmented T1 images of 15 AVH patients with SimNIBS 3.2. Then we utilized the head surface nodes of these patients to generate their individual parameter space (Jiang et al., 2022). We utilized the projection method in Anatomist software (Andoh et al., 2009). Specifically, the MNI coordinates of individual stimulation locations were converted to subject space with the mni2subject_coords function of the SimNIBS 3.2 package and then projected on the scalp by calculating the center of gravity of the (n = 10) closest head surface nodes. Coil orientation was quantified according to the recorded stimulation parameter (Paillère-Martinot et al., 2017). We first defined a normal vector that was averaged over a round area around the target site (5 mm of radius). The handle direction was perpendicular to the plane through the TP3 line and the normal vector of the scalp target site. If the target was on the right hemisphere, the handle direction was perpendicular to the plane through the TP4 line and the normal vector of the scalp target site (Figure 4A). In addition, the coil placements of the cohort are also shown in parameter space (Supplementary Table 6).

##### 1.2.2. Electric field modeling

Isotropic tissue conductivities were assigned as $\sigma_{scalp}$ = 0.465 S/m, $\sigma_{bone}$ = 0.010 S/m, $\sigma_{CSF}$ = 1.654 S/m, $\sigma_{GM}$ = 0.275 S/m, and $\sigma_{WM}$ = 0.126 S/m (Wagner et al., 2004). Magstim 70 mm figure-of-8 coil was chosen for electric field simulations, the same as that in actual treatment (Thielscher and Kammer, 2004; Paillère Martinot et al., 2010; Paillère-Martinot et al., 2017). The electric field was generated with the actual coil placement with SimNIBS 3.2. We took Gray matter voxels with E-field >90% of the maximum peak value (Makarov et al., 2021) as the stimulated region by TMS. To avoid the effect of outliers, the peak value is defined as the 99.9th percentile of maximal E-field (Saturnino et al., 2019).

The kept E-field map was down-sampled from 1 mm × 1 mm × 1 mm to 3 mm × 3 mm × 3 mm and masked with the former gray-matter mask. According to equation 1, a weight was assigned to each voxel by normalizing the suprathreshold E-field to sum 1.

##### 1.2.3. Group-level functional connectivity

The rsfMRI data from the SLIM dataset were preprocessed with the DPABI toolbox (Yan et al., 2016), which included the following steps: 1) elimination of the first ten time points; 2) correction for slice timing; 3) realignment of the functional image to correct for head motion; 4) regression of nuisance signals estimated from the signals of white matter, CSF and the mean global signal of gray matter (Fox et al., 2012); 5) 0.01~0.1Hz band-pass filtering, and 6) spatial smoothing (kernel FWHM 6 mm × 6 mm × 6 mm). The functional images were co-registered to scalp-extracted anatomical images and then normalized into MNI space with the DARTEL algorithm (Ashburner, 2007).

In NMI space, the spatially normalized functional images were masked with a customized gray-matter mask. The gray-matter mask was produced from the Tissue Probability Atlas provided by SPM, excluding voxels with a probability of less than 20%. Then the masked image was resampled into 3 mm × 3 mm × 3 mm spacing to be consistent with the functional images. As a result, the final gray matter images consisted of 50740 voxels in MNI space, the same across participants. Finally, a voxel-to-voxel correlation matrix (50740×50740) was calculated for each participant using Pearson's correlation. Then these matrices of 512 subjects were averaged to generate the group-level connectivity matrix.

##### *1.2.4. TMS stimulation network*

According to equation 2, the network-level effect was computed from the individual E-field weight and group-level voxel-to-voxel functional connectivity in MNI space.

##### *1.2.5. Pathological network from meta-analysis*

Results from one coordinate-based meta-analysis (CBMA) study were utilized based on descriptions of the pathological network for MDD in the network targeting model (Gray et al., 2020). In total, 3594 MDD patients, and 3362 healthy controls were included in the meta-analysis. In this study, voxel-based pathophysiology changes were compared between patients with MDD and healthy controls. The hyper-activation is defined as the increased function relative to controls. The hypo-activation is defined as the decreased function relative to controls. The resultant map of the meta-analysis was produced via the ALE technique, and foci of significant hyper- and hypo-activity were reported in the Talairach space. Specifically, 6 hyper-activated and 3 hypo-activated foci are listed in the ALE results (Supplementary Table 1). All coordinates were transformed using the tal2icbm transformation (Laird et al., 2009). The radius of foci was set to 10 mm (Fox et al., 2013; Cash et al., 2019).

Results from one CBMA study were utilized based on descriptions of the pathological network for AVH in the network targeting model (Kühn and Gallinat, 2012). In this study, regional activities were compared between patients with AVH and healthy controls in two types of studies. Specifically, one type focused on comparing the brain states of AVH patients during periods with and without hallucination. The other type focused on comparing the brain states of hallucinating schizophrenic patients versus non-hallucinating schizophrenic patients or healthy controls. In total, 166 hallucinators, 39 non-hallucinators, and 69 healthy controls were included in the meta-analysis. The resultant map of the meta-analysis was produced via the ALE technique, and foci of significant hyper- and hypo-activity were reported in the Talairach space. Specifically, 4 hyper-activated and 6 hypo-activated foci are listed in the ALE results (Supplementary Table 2). All of the coordinates were transformed using the tal2icbm transformation (Laird et al., 2009). The radius of foci was set to 10 mm (Fox et al., 2013; Cash et al., 2019).

##### *1.2.6. Network targeting accuracy on pathological network*

According to equation 5, the network targeting accuracy was calculated by spatially anti-correlating the TMS stimulation network and pathological network derived from meta-analysis results for MDD or AVH.

#### 1.3. **Validation of network targeting models**

In the first validation experiment on depression, to test the hypothesis that NTA can predict clinical improvement, we conducted a correlation analysis between the averaged NTA across subjects for the DLPFC target and the expected clinical improvement of HDRS of each DLPFC target estimated by Herbsman's equation (Herbsman et al., 2009): HDRS drop = -.84 + (X * -.022) + (Y * .012).

In the second validation experiment on MDD, to test the hypothesis that NTA can predict clinical improvement, we conducted a correlation analysis between NTA of the left PFC group and clinical improvement of MADRS. Clinical outcome was measured by the proportional decrease of total score between pre-treatment and post-treatment assessments. In addition, we conducted a correlation analysis between the NTA of the right PFC group and the clinical improvement of MADRS (Figure S6). We conducted a correlation analysis between the NTA of the double-side PFC group and the clinical improvement of MADRS (Figure S7).

In the third validation experiment on AVH, to test the hypothesis that NTA can predict clinical improvement, we conducted a correlation analysis between NTA and clinical improvement of AHRS. Clinical outcome was measured by the proportional decrease of total score between pre-treatment and post-treatment assessments. In further exploration of symptom specificity, we conducted a correlation analysis between NTA and clinical improvement of SAPS, and we conducted a correlation analysis between NTA and clinical improvement of SANS.

#### 1.4. **Control Analysis**

##### *1.4.1. Permutation Tests*

To confirm our hypothesis, we conducted two permutation tests in all validation experiments. To test whether the NTA based on the pathological network can predict the treatment outcome better than that based on a random network, we generated pseudo networks by randomly permuting the spatial distribution of hyper- and hypo-activated foci of the pathological network 10,000 times. NTA derived from the pseudo networks was used to predict the treatment outcome. The predictiveness values were taken as the null distribution (Figure S2). To test the dependency of NTA and treatment outcome in a non-parametric way, we randomly reassigned clinical outcomes among patients, repeating 10,000 times. The null distribution of predictiveness was obtained by correlating NTA with these permutated clinical outcomes (Figure S3).

##### *1.4.2. Robustness Tests*

The E-field threshold is an unsolved factor that may affect network targeting accuracy. To test the robustness of our results, we calculated NTA with different thresholds (75%, 80%, 85%, 95%, 99%) in all validation experiments. Then we calculated the similarity between the NTA of the current threshold and the NTAs of other thresholds (Opitz et al., 2016) in all experiments (Figure S4).

In the construction of the pathological network from meta-analysis, we created concentric spheres to estimate the abnormal foci. We used a radius of 10 mm referring to a previous study (Fox et al., 2013). In addition, we created spheres with other foci radii (4, 6, 8, 12, 16 mm) and calculated the NTAs in all validation experiments. We compared the NTAs derived from different sphere radii along the participants (Figure S5).

##### *1.4.3. Comparing network targeting to the SGC-FC method*

We compared network targeting with SGC-FC in the MDD cohort in the first and second validation experiments. The SGC-FC method (Fox et al., 2012) targets one deep node, the SGC. As an alternative method, the network targeting method targets multiple nodes of the MDD pathological network. To compare these methods, we calculate the targeting score with the SGC method. More details are shown in Figure S8.

##### *1.4.4. Comparing network targeting to location-based targeting*

In the third validation experiment, as positive controls, we compared the proposed NTA model to the traditional coordinates-based targeting model used for AVH treatment. Traditionally, TP3 (Hoffman et al., 2003) and L.Wernicke (Hoffman et al., 2013) are selected as stimulation targets to set TMS coils, assuming that coverage of the particular ROI is related to treatment outcome. In this assumption, targeting accuracy to the ROI can be quantified with an index of coordinate-based targeting score (CTS), defined as the reciprocal of the distance between the actual placement and traditional target. CTS is hypothesized to predict clinical improvement (Figure S10).

Specifically, cortical sites were [-57, -49, 28] for TP3 (Herwig et al., 2003) and [-65, -41, 9] for L.Wernicke (Hoffman et al., 2013). These coordinates were transformed to MNI space with tal2icbm (Laird et al., 2009). Since the traditional target site is in the left hemisphere, one patient treated in the right hemisphere was excluded from the current analysis.

### 2. Individualized parameter optimization

Having been validated to predict treatment efficacy from the stimulation parameter, NTA in principle is capable to identify the TMS coil placement with an optimal outcome. Here, on the two clinical cohorts of MDD (27 patients, Figure 3) and AVH (15 patients, Figure 4) received active TMS treatments, we simulated coil settings for a range of the parameter space. We then examined the variation across individuals in the optimal coil placement.

#### 2.1. Searching space of the stimulation parameter

Position: In the first MDD simulation experiment, a cranial search space that included 125 CPC positions and covered a broad area around the left DLPFC (Lefaucheur et al., 2014; Xiao et al., 2018; Cash et al., 2020; Balderston et al., 2021), was chosen for the experiment (Figure 5). The CPC positions ($p_{NZ}$, $p_{AL}$) were the same across the subjects (Supplementary Table 7). In the second AVH simulation experiment, a cranial search space that included 122 CPC positions and covered a broad area including left STG and left TPJ, ROIs adopted in TMS treatments for AVH (Hoffman et al., 2003, 2013; Klirova et al., 2013; Lefaucheur et al., 2014; Paillère-Martinot et al., 2017; Xiao et al., 2018), was chosen for the experiment. The CPC positions ($p_{NZ}$, $p_{AL}$) were the same across the subjects (Supplementary Table 8). The distance between two adjacent CPC positions on the individual head model was around 6.5 mm.

Orientation: In all simulation experiments, the coil was only rotated around its yaw axis and fixed on its roll axis and pitch axis. For each position, 12 different orientations (0°, -15°, -30°, -45°, -60°, -75°, -90°, -105°, -120°, -135°, -150°, -165°, 15-degree intervals) were investigated.

#### 2.2. Position-by-orientation interaction of the NTA

Network targeting accuracy was calculated for each pair of position and orientation. In the first MDD simulation experiment, a total of 1500 NTA (125 positions × 12 orientations) were calculated for each individual. In the second AVH simulation experiment, a total of 1464 NTA (122 positions × 12 orientations) were calculated for each individual. In all simulation experiments, an analysis of variance (ANOVA) was used to test for NTA differences among coil positions and coil orientations.

For visualization purposes, in the first MDD simulation experiment, we interpolated NTA search space from 125×12 to 27977 × 12 using the natural neighbor interpolation method (Amidror, 2002), as shown in Figure 5. In the second AVH simulation experiment, we interpolated NTA search space from 122×12 to 58470 × 12 using natural interpolation (Figure 6). The optimal TMS parameters were the positions and orientations leading to maximum NTA (Figure 5 & Figure 6 & Supplementary Table 9 & Supplementary Table 10).

### Supplementary Results

**Supplementary Table 1. The foci coordinates of MDD meta-analysis** (Gray et al., 2020)**.**

**Supplementary Table 2. The foci coordinates of AVH meta-analysis** (Kühn and Gallinat, 2012)**.**

**Supplementary Table 3. The coil placements of the empirical DLPFC cohort are shown in parameter space (Part A).**

**Supplementary Table 4. The coil placements of the empirical DLPFC cohort are shown in parameter space (Part B).**

**Supplementary Table 5. The coil placements of the clinical MDD cohort are shown in parameter space.**

**Supplementary Table 6. The coil placements of the clinical AVH cohort are shown in parameter space.**

**Supplementary Table 7. CPC positions of MDD simulation experiment.**

**Supplementary Table 8. CPC positions of AVH simulation experiment.**

**Supplementary Table 9. The optimal TMS coil placements of MDD simulation experiment.**

**Supplementary Table 10. The optimal TMS coil placements of AVH simulation experiment.**

**Supplementary Figure 1. Steps of finding a scalp position based on a DLPFC site.**

**Supplementary Figure 2. Permutation tests of targeting pseudo networks for assessing the predictiveness of NTA.**

**Supplementary Figure 3. Permutation tests of randomized prediction for assessing the predictiveness of NTA.**

**Supplementary Figure 4. Robustness of E-field threshold.**

**Supplementary Figure 5. Robustness of radii of foci of the pathological network.**

**Supplementary Figure 6. Network targeting accuracy failed to predict treatment efficacy in the clinical MDD cohort who received TMS in the right PFC.**

**Supplementary Figure 7. Network targeting accuracy failed to predict treatment efficacy in the clinical MDD cohort who received TMS on both sides of PFC.**

**Supplementary Figure 8. Comparing network targeting with the SGC-FC method in depression cohorts.**

**Supplementary Figure 9. Coordinates-based targeting method in AVH treatment.**

**Supplementary Table 1. The foci coordinates of MDD meta-analysis** (Gray et al., 2020)**.**

| Foci Name | Talairach Coordinates | | | Activation |
|---|---|---|---|---|
| | x | y | z | |
| Hippocampus | -24 | -16 | -15 | **Hyper-** |
| Subgenual cingulate | 2 | 30 | -2 | **Hyper-** |
| Amygdala/putamen | 21 | 2 | -9 | **Hyper-** |
| Hippocampus | -24 | -17 | -15 | **Hyper-** |
| Amygdala/putamen | 20 | 4 | -9 | **Hyper-** |
| Hippocampus | -25 | -17 | -15 | **Hyper-** |
| Middle occipital/inferior temporal gyrus | 41 | -66 | 1 | Hypo- |
| Retrosplenial cortex | -12 | -47 | 4 | Hypo- |
| Putamen | 26 | 6 | 10 | Hypo- |

**Supplementary Table 2. The foci coordinates of AVH meta-analysis** (Kühn and Gallinat, 2012)**.**

| **Foci Name** | **Talairach Coordinates** | | | **Activation** |
|---|---|---|---|---|
| | **x** | **y** | **z** | |
| Left parietal operculum | -55 | -19 | 16 | **Hyper-** |
| Left postcentral gyrus | -49 | -17 | 41 | **Hyper-** |
| Right postcentral gyrus | 36 | -32 | 50 | **Hyper-** |
| Left inferior frontal gyrus | -48 | 2 | 6 | **Hyper-** |
| Left middle temporal gyrus | -56 | -30 | 0 | Hypo- |
| Left premotor cortex | -10 | 3 | 56 | Hypo- |
| Anterior cingulate gyrus | -44 | -22 | 0 | Hypo- |
| Left superior temporal gyrus | -42 | 2 | 18 | Hypo- |
| Anterior cingulate gyrus | -9 | 4 | 37 | Hypo- |
| Anterior cingulate gyrus | -4 | 26 | 31 | Hypo- |

#### Supplementary Table 3. The coil placements of the empirical DLPFC cohort are shown in parameter space (Part A).

| Sub ID | 5cm | | 5.5cm | | Nonresponder | | Responder | | Fitzgerald | | Rusjan | | Ori |
|---|---|---|---|---|---|---|---|---|---|---|---|---|---|
| | Position ($s$) | | Position ($s$) | | Position ($s$) | | Position ($s$) | | Position ($s$) | | Position ($s$) | | ($\theta$) |
| | $p_{NZ}$ | $p_{AL}$ | $p_{NZ}$ | $p_{AL}$ | $p_{NZ}$ | $p_{AL}$ | $p_{NZ}$ | $p_{AL}$ | $p_{NZ}$ | $p_{AL}$ | $p_{NZ}$ | $p_{AL}$ | |
| 1 | 0.365 | 0.325 | 0.345 | 0.370 | 0.365 | 0.325 | 0.365 | 0.325 | 0.280 | 0.330 | 0.320 | 0.300 | -45° |
| 2 | 0.365 | 0.335 | 0.335 | 0.370 | 0.365 | 0.335 | 0.355 | 0.325 | 0.270 | 0.335 | 0.310 | 0.300 | -45° |
| 3 | 0.355 | 0.330 | 0.335 | 0.365 | 0.355 | 0.330 | 0.350 | 0.325 | 0.265 | 0.330 | 0.305 | 0.300 | -45° |
| 4 | 0.345 | 0.335 | 0.335 | 0.365 | 0.345 | 0.340 | 0.345 | 0.325 | 0.265 | 0.330 | 0.305 | 0.300 | -45° |
| 5 | 0.370 | 0.320 | 0.345 | 0.360 | 0.370 | 0.320 | 0.365 | 0.315 | 0.280 | 0.320 | 0.320 | 0.290 | -45° |
| 6 | 0.350 | 0.335 | 0.335 | 0.365 | 0.350 | 0.340 | 0.350 | 0.325 | 0.270 | 0.330 | 0.305 | 0.305 | -45° |
| 7 | 0.350 | 0.335 | 0.335 | 0.365 | 0.350 | 0.335 | 0.350 | 0.320 | 0.265 | 0.330 | 0.310 | 0.295 | -45° |
| 8 | 0.365 | 0.325 | 0.340 | 0.350 | 0.365 | 0.325 | 0.360 | 0.310 | 0.270 | 0.325 | 0.315 | 0.285 | -45° |
| 9 | 0.350 | 0.335 | 0.335 | 0.365 | 0.350 | 0.340 | 0.350 | 0.325 | 0.270 | 0.330 | 0.310 | 0.295 | -45° |
| 10 | 0.360 | 0.335 | 0.340 | 0.365 | 0.360 | 0.335 | 0.360 | 0.320 | 0.270 | 0.330 | 0.310 | 0.295 | -45° |
| 11 | 0.370 | 0.340 | 0.345 | 0.365 | 0.370 | 0.340 | 0.365 | 0.325 | 0.280 | 0.335 | 0.320 | 0.300 | -45° |
| 12 | 0.355 | 0.340 | 0.335 | 0.370 | 0.355 | 0.340 | 0.355 | 0.325 | 0.270 | 0.335 | 0.310 | 0.300 | -45° |
| 13 | 0.350 | 0.340 | 0.330 | 0.365 | 0.350 | 0.340 | 0.350 | 0.325 | 0.265 | 0.335 | 0.305 | 0.300 | -45° |
| 14 | 0.365 | 0.325 | 0.340 | 0.360 | 0.365 | 0.325 | 0.365 | 0.320 | 0.275 | 0.330 | 0.315 | 0.295 | -45° |
| 15 | 0.360 | 0.340 | 0.335 | 0.370 | 0.360 | 0.340 | 0.355 | 0.325 | 0.275 | 0.335 | 0.310 | 0.300 | -45° |
| 16 | 0.350 | 0.335 | 0.330 | 0.370 | 0.350 | 0.335 | 0.350 | 0.325 | 0.265 | 0.335 | 0.305 | 0.300 | -45° |
| 17 | 0.360 | 0.335 | 0.335 | 0.365 | 0.360 | 0.340 | 0.360 | 0.320 | 0.270 | 0.330 | 0.310 | 0.295 | -45° |
| 18 | 0.355 | 0.335 | 0.330 | 0.365 | 0.355 | 0.340 | 0.355 | 0.320 | 0.270 | 0.330 | 0.310 | 0.300 | -45° |
| 19 | 0.355 | 0.340 | 0.340 | 0.370 | 0.355 | 0.340 | 0.355 | 0.325 | 0.275 | 0.330 | 0.310 | 0.300 | -45° |
| 20 | 0.355 | 0.335 | 0.335 | 0.365 | 0.355 | 0.340 | 0.355 | 0.325 | 0.270 | 0.335 | 0.310 | 0.300 | -45° |
| 21 | 0.350 | 0.340 | 0.335 | 0.370 | 0.350 | 0.345 | 0.350 | 0.330 | 0.270 | 0.335 | 0.310 | 0.305 | -45° |
| 22 | 0.355 | 0.335 | 0.335 | 0.370 | 0.355 | 0.335 | 0.355 | 0.330 | 0.275 | 0.335 | 0.310 | 0.305 | -45° |
| 23 | 0.345 | 0.325 | 0.330 | 0.360 | 0.345 | 0.330 | 0.345 | 0.315 | 0.270 | 0.325 | 0.305 | 0.290 | -45° |
| 24 | 0.355 | 0.330 | 0.335 | 0.360 | 0.355 | 0.335 | 0.355 | 0.320 | 0.270 | 0.330 | 0.310 | 0.295 | -45° |
| 25 | 0.370 | 0.340 | 0.345 | 0.370 | 0.370 | 0.340 | 0.365 | 0.325 | 0.280 | 0.330 | 0.320 | 0.300 | -45° |
| 26 | 0.360 | 0.335 | 0.335 | 0.365 | 0.360 | 0.335 | 0.360 | 0.320 | 0.275 | 0.330 | 0.315 | 0.300 | -45° |
| 27 | 0.365 | 0.330 | 0.340 | 0.365 | 0.365 | 0.330 | 0.365 | 0.325 | 0.285 | 0.330 | 0.320 | 0.300 | -45° |
| 28 | 0.350 | 0.330 | 0.330 | 0.365 | 0.350 | 0.335 | 0.350 | 0.320 | 0.260 | 0.330 | 0.300 | 0.290 | -45° |
| 29 | 0.370 | 0.330 | 0.350 | 0.360 | 0.370 | 0.335 | 0.370 | 0.320 | 0.280 | 0.325 | 0.320 | 0.290 | -45° |
| 30 | 0.365 | 0.330 | 0.335 | 0.365 | 0.365 | 0.330 | 0.360 | 0.315 | 0.275 | 0.335 | 0.315 | 0.295 | -45° |
| 31 | 0.350 | 0.335 | 0.335 | 0.370 | 0.350 | 0.340 | 0.350 | 0.320 | 0.270 | 0.330 | 0.305 | 0.295 | -45° |
| 32 | 0.360 | 0.335 | 0.335 | 0.370 | 0.360 | 0.335 | 0.355 | 0.325 | 0.275 | 0.330 | 0.310 | 0.300 | -45° |
| 33 | 0.350 | 0.330 | 0.330 | 0.370 | 0.350 | 0.330 | 0.350 | 0.325 | 0.265 | 0.330 | 0.300 | 0.300 | -45° |
| 34 | 0.345 | 0.340 | 0.325 | 0.370 | 0.345 | 0.340 | 0.345 | 0.325 | 0.265 | 0.335 | 0.300 | 0.305 | -45° |
| 35 | 0.350 | 0.340 | 0.335 | 0.370 | 0.350 | 0.345 | 0.350 | 0.325 | 0.265 | 0.335 | 0.305 | 0.300 | -45° |
| 36 | 0.360 | 0.330 | 0.345 | 0.365 | 0.360 | 0.330 | 0.360 | 0.320 | 0.280 | 0.325 | 0.320 | 0.290 | -45° |
| 37 | 0.355 | 0.335 | 0.335 | 0.370 | 0.355 | 0.335 | 0.355 | 0.325 | 0.270 | 0.335 | 0.310 | 0.300 | -45° |
| 38 | 0.360 | 0.335 | 0.340 | 0.365 | 0.360 | 0.335 | 0.360 | 0.320 | 0.280 | 0.330 | 0.320 | 0.295 | -45° |
| 39 | 0.355 | 0.335 | 0.335 | 0.365 | 0.355 | 0.340 | 0.355 | 0.320 | 0.270 | 0.330 | 0.315 | 0.295 | -45° |
| 40 | 0.360 | 0.340 | 0.335 | 0.370 | 0.360 | 0.345 | 0.355 | 0.325 | 0.270 | 0.335 | 0.310 | 0.300 | -45° |
| 41 | 0.365 | 0.320 | 0.345 | 0.365 | 0.360 | 0.340 | 0.365 | 0.320 | 0.280 | 0.330 | 0.315 | 0.295 | -45° |
| 42 | 0.360 | 0.330 | 0.340 | 0.365 | 0.360 | 0.330 | 0.360 | 0.320 | 0.275 | 0.325 | 0.310 | 0.295 | -45° |
| 43 | 0.350 | 0.335 | 0.330 | 0.375 | 0.350 | 0.335 | 0.350 | 0.330 | 0.265 | 0.340 | 0.310 | 0.305 | -45° |
| 44 | 0.355 | 0.335 | 0.335 | 0.360 | 0.355 | 0.335 | 0.355 | 0.320 | 0.270 | 0.325 | 0.310 | 0.295 | -45° |
| 45 | 0.350 | 0.330 | 0.335 | 0.360 | 0.350 | 0.335 | 0.350 | 0.320 | 0.270 | 0.325 | 0.310 | 0.295 | -45° |
| 46 | 0.370 | 0.330 | 0.345 | 0.365 | 0.370 | 0.330 | 0.365 | 0.325 | 0.280 | 0.330 | 0.320 | 0.295 | -45° |
| 47 | 0.350 | 0.340 | 0.335 | 0.370 | 0.350 | 0.340 | 0.350 | 0.325 | 0.265 | 0.335 | 0.305 | 0.305 | -45° |
| 48 | 0.340 | 0.335 | 0.330 | 0.365 | 0.340 | 0.335 | 0.340 | 0.325 | 0.265 | 0.335 | 0.300 | 0.300 | -45° |
| 49 | 0.360 | 0.330 | 0.335 | 0.370 | 0.360 | 0.330 | 0.355 | 0.330 | 0.270 | 0.335 | 0.310 | 0.300 | -45° |
| 50 | 0.355 | 0.335 | 0.335 | 0.360 | 0.355 | 0.335 | 0.355 | 0.320 | 0.275 | 0.330 | 0.315 | 0.300 | -45° |
| 51 | 0.355 | 0.335 | 0.340 | 0.365 | 0.355 | 0.335 | 0.355 | 0.325 | 0.270 | 0.330 | 0.310 | 0.300 | -45° |
| 52 | 0.355 | 0.335 | 0.340 | 0.370 | 0.355 | 0.335 | 0.355 | 0.325 | 0.275 | 0.340 | 0.315 | 0.300 | -45° |
| 53 | 0.360 | 0.330 | 0.335 | 0.365 | 0.360 | 0.330 | 0.355 | 0.325 | 0.270 | 0.330 | 0.310 | 0.300 | -45° |
| 54 | 0.345 | 0.330 | 0.330 | 0.370 | 0.345 | 0.330 | 0.345 | 0.325 | 0.265 | 0.330 | 0.300 | 0.300 | -45° |
| 55 | 0.355 | 0.335 | 0.330 | 0.360 | 0.355 | 0.335 | 0.355 | 0.320 | 0.270 | 0.330 | 0.310 | 0.295 | -45° |
| 56 | 0.350 | 0.335 | 0.335 | 0.370 | 0.350 | 0.335 | 0.350 | 0.325 | 0.270 | 0.335 | 0.310 | 0.300 | -45° |
| 57 | 0.365 | 0.335 | 0.330 | 0.360 | 0.365 | 0.335 | 0.350 | 0.320 | 0.270 | 0.330 | 0.305 | 0.295 | -45° |
| 58 | 0.350 | 0.335 | 0.330 | 0.370 | 0.350 | 0.335 | 0.350 | 0.330 | 0.265 | 0.340 | 0.305 | 0.300 | -45° |
| 59 | 0.365 | 0.340 | 0.335 | 0.365 | 0.365 | 0.340 | 0.360 | 0.325 | 0.270 | 0.335 | 0.310 | 0.300 | -45° |
| 60 | 0.350 | 0.335 | 0.325 | 0.370 | 0.350 | 0.335 | 0.350 | 0.325 | 0.265 | 0.335 | 0.300 | 0.300 | -45° |
| 61 | 0.345 | 0.340 | 0.335 | 0.365 | 0.345 | 0.340 | 0.345 | 0.325 | 0.265 | 0.330 | 0.305 | 0.295 | -45° |
| 62 | 0.345 | 0.350 | 0.335 | 0.370 | 0.345 | 0.350 | 0.350 | 0.330 | 0.265 | 0.340 | 0.310 | 0.305 | -45° |
| 63 | 0.360 | 0.330 | 0.340 | 0.370 | 0.360 | 0.330 | 0.360 | 0.325 | 0.275 | 0.330 | 0.310 | 0.295 | -45° |
| 64 | 0.360 | 0.335 | 0.340 | 0.370 | 0.360 | 0.340 | 0.360 | 0.325 | 0.275 | 0.335 | 0.315 | 0.300 | -45° |
| 65 | 0.360 | 0.335 | 0.335 | 0.365 | 0.360 | 0.335 | 0.355 | 0.325 | 0.275 | 0.330 | 0.315 | 0.300 | -45° |

| | | | | | | | | | | | | | |
|---|---|---|---|---|---|---|---|---|---|---|---|---|---|
| 66 | 0.350 | 0.340 | 0.335 | 0.365 | 0.350 | 0.340 | 0.350 | 0.325 | 0.270 | 0.330 | 0.310 | 0.300 | -45° |
| 67 | 0.365 | 0.335 | 0.335 | 0.365 | 0.365 | 0.335 | 0.360 | 0.325 | 0.275 | 0.330 | 0.310 | 0.300 | -45° |
| 68 | 0.360 | 0.330 | 0.335 | 0.365 | 0.360 | 0.330 | 0.355 | 0.325 | 0.270 | 0.330 | 0.310 | 0.300 | -45° |

#### Supplementary Table 4. The coil placements of the empirical DLPFC cohort are shown in parameter space (Part B).

| Sub ID | Paus/Cho Position ($s$) | | BA9 Position ($s$) | | BA46 Position ($s$) | | F3 Herwig Position ($s$) | | F3 Okamoto Position ($s$) | | Beam F3 Position ($s$) | | Ori |
|---|---|---|---|---|---|---|---|---|---|---|---|---|---|
| | $p_{NZ}$ | $p_{AL}$ | $p_{NZ}$ | $p_{AL}$ | $p_{NZ}$ | $p_{AL}$ | $p_{NZ}$ | $p_{AL}$ | $p_{NZ}$ | $p_{AL}$ | $p_{NZ}$ | $p_{AL}$ | ($\theta$) |
| 1 | 0.305 | 0.320 | 0.305 | 0.355 | 0.270 | 0.315 | 0.355 | 0.350 | 0.255 | 0.360 | 0.265 | 0.330 | -45° |
| 2 | 0.300 | 0.320 | 0.295 | 0.360 | 0.260 | 0.320 | 0.345 | 0.350 | 0.250 | 0.365 | 0.255 | 0.335 | -45° |
| 3 | 0.295 | 0.320 | 0.295 | 0.355 | 0.260 | 0.315 | 0.340 | 0.350 | 0.240 | 0.360 | 0.250 | 0.330 | -45° |
| 4 | 0.295 | 0.315 | 0.295 | 0.355 | 0.255 | 0.310 | 0.340 | 0.350 | 0.240 | 0.360 | 0.250 | 0.330 | -45° |
| 5 | 0.310 | 0.305 | 0.305 | 0.345 | 0.270 | 0.305 | 0.355 | 0.340 | 0.255 | 0.350 | 0.260 | 0.320 | -45° |
| 6 | 0.300 | 0.320 | 0.300 | 0.355 | 0.260 | 0.315 | 0.345 | 0.350 | 0.245 | 0.360 | 0.255 | 0.330 | -45° |
| 7 | 0.300 | 0.310 | 0.295 | 0.355 | 0.260 | 0.310 | 0.345 | 0.350 | 0.240 | 0.365 | 0.250 | 0.330 | -45° |
| 8 | 0.310 | 0.305 | 0.300 | 0.340 | 0.265 | 0.305 | 0.350 | 0.335 | 0.240 | 0.355 | 0.255 | 0.325 | -45° |
| 9 | 0.305 | 0.315 | 0.295 | 0.355 | 0.265 | 0.310 | 0.345 | 0.350 | 0.245 | 0.360 | 0.255 | 0.330 | -45° |
| 10 | 0.300 | 0.315 | 0.300 | 0.355 | 0.260 | 0.315 | 0.350 | 0.350 | 0.245 | 0.360 | 0.255 | 0.330 | -45° |
| 11 | 0.310 | 0.315 | 0.305 | 0.355 | 0.270 | 0.310 | 0.355 | 0.350 | 0.255 | 0.360 | 0.265 | 0.330 | -45° |
| 12 | 0.300 | 0.320 | 0.295 | 0.360 | 0.255 | 0.315 | 0.345 | 0.350 | 0.245 | 0.360 | 0.250 | 0.330 | -45° |
| 13 | 0.295 | 0.320 | 0.290 | 0.355 | 0.260 | 0.315 | 0.340 | 0.350 | 0.240 | 0.365 | 0.250 | 0.335 | -45° |
| 14 | 0.305 | 0.310 | 0.300 | 0.355 | 0.265 | 0.310 | 0.350 | 0.345 | 0.250 | 0.360 | 0.260 | 0.330 | -45° |
| 15 | 0.300 | 0.320 | 0.300 | 0.360 | 0.265 | 0.315 | 0.345 | 0.350 | 0.245 | 0.365 | 0.255 | 0.335 | -45° |
| 16 | 0.295 | 0.315 | 0.290 | 0.355 | 0.260 | 0.315 | 0.340 | 0.350 | 0.240 | 0.365 | 0.250 | 0.335 | -45° |
| 17 | 0.305 | 0.315 | 0.300 | 0.355 | 0.265 | 0.315 | 0.350 | 0.345 | 0.245 | 0.360 | 0.255 | 0.330 | -45° |
| 18 | 0.300 | 0.320 | 0.295 | 0.355 | 0.260 | 0.315 | 0.340 | 0.345 | 0.245 | 0.355 | 0.255 | 0.330 | -45° |
| 19 | 0.305 | 0.325 | 0.300 | 0.360 | 0.265 | 0.315 | 0.350 | 0.355 | 0.250 | 0.355 | 0.255 | 0.330 | -45° |
| 20 | 0.305 | 0.315 | 0.300 | 0.355 | 0.265 | 0.315 | 0.345 | 0.350 | 0.245 | 0.365 | 0.255 | 0.335 | -45° |
| 21 | 0.305 | 0.325 | 0.300 | 0.360 | 0.265 | 0.320 | 0.345 | 0.355 | 0.245 | 0.360 | 0.255 | 0.335 | -45° |
| 22 | 0.300 | 0.330 | 0.295 | 0.360 | 0.265 | 0.320 | 0.345 | 0.350 | 0.250 | 0.360 | 0.255 | 0.335 | -45° |
| 23 | 0.295 | 0.310 | 0.295 | 0.350 | 0.260 | 0.310 | 0.340 | 0.340 | 0.240 | 0.355 | 0.250 | 0.325 | -45° |
| 24 | 0.305 | 0.310 | 0.300 | 0.350 | 0.265 | 0.310 | 0.350 | 0.345 | 0.245 | 0.355 | 0.255 | 0.325 | -45° |
| 25 | 0.310 | 0.320 | 0.305 | 0.360 | 0.275 | 0.315 | 0.355 | 0.355 | 0.250 | 0.360 | 0.265 | 0.330 | -45° |
| 26 | 0.300 | 0.320 | 0.295 | 0.355 | 0.265 | 0.315 | 0.345 | 0.345 | 0.250 | 0.360 | 0.255 | 0.335 | -45° |
| 27 | 0.305 | 0.310 | 0.305 | 0.350 | 0.275 | 0.310 | 0.350 | 0.345 | 0.260 | 0.360 | 0.265 | 0.330 | -45° |
| 28 | 0.290 | 0.310 | 0.290 | 0.355 | 0.255 | 0.310 | 0.340 | 0.345 | 0.235 | 0.360 | 0.245 | 0.330 | -45° |
| 29 | 0.305 | 0.305 | 0.300 | 0.345 | 0.270 | 0.305 | 0.365 | 0.345 | 0.250 | 0.355 | 0.260 | 0.325 | -45° |
| 30 | 0.305 | 0.310 | 0.300 | 0.360 | 0.265 | 0.315 | 0.345 | 0.340 | 0.255 | 0.365 | 0.255 | 0.335 | -45° |
| 31 | 0.290 | 0.315 | 0.295 | 0.360 | 0.255 | 0.310 | 0.345 | 0.350 | 0.245 | 0.360 | 0.250 | 0.330 | -45° |
| 32 | 0.300 | 0.320 | 0.300 | 0.355 | 0.265 | 0.315 | 0.345 | 0.350 | 0.245 | 0.360 | 0.255 | 0.330 | -45° |
| 33 | 0.285 | 0.315 | 0.290 | 0.355 | 0.250 | 0.315 | 0.340 | 0.350 | 0.240 | 0.360 | 0.245 | 0.330 | -45° |
| 34 | 0.290 | 0.320 | 0.290 | 0.360 | 0.255 | 0.315 | 0.335 | 0.350 | 0.235 | 0.365 | 0.245 | 0.335 | -45° |
| 35 | 0.295 | 0.320 | 0.290 | 0.360 | 0.260 | 0.315 | 0.345 | 0.355 | 0.240 | 0.360 | 0.250 | 0.335 | -45° |
| 36 | 0.305 | 0.310 | 0.305 | 0.355 | 0.265 | 0.305 | 0.355 | 0.345 | 0.250 | 0.355 | 0.260 | 0.325 | -45° |
| 37 | 0.300 | 0.320 | 0.295 | 0.360 | 0.265 | 0.315 | 0.345 | 0.350 | 0.250 | 0.365 | 0.255 | 0.335 | -45° |
| 38 | 0.310 | 0.315 | 0.305 | 0.355 | 0.270 | 0.310 | 0.350 | 0.345 | 0.250 | 0.355 | 0.260 | 0.330 | -45° |
| 39 | 0.305 | 0.315 | 0.295 | 0.355 | 0.265 | 0.310 | 0.350 | 0.350 | 0.240 | 0.355 | 0.255 | 0.330 | -45° |
| 40 | 0.300 | 0.320 | 0.300 | 0.360 | 0.260 | 0.315 | 0.345 | 0.355 | 0.245 | 0.360 | 0.255 | 0.335 | -45° |
| 41 | 0.310 | 0.315 | 0.305 | 0.355 | 0.270 | 0.310 | 0.355 | 0.350 | 0.250 | 0.360 | 0.260 | 0.330 | -45° |
| 42 | 0.300 | 0.315 | 0.300 | 0.350 | 0.265 | 0.310 | 0.350 | 0.350 | 0.250 | 0.355 | 0.260 | 0.325 | -45° |
| 43 | 0.300 | 0.325 | 0.295 | 0.365 | 0.260 | 0.320 | 0.345 | 0.355 | 0.240 | 0.365 | 0.250 | 0.340 | -45° |
| 44 | 0.300 | 0.315 | 0.295 | 0.350 | 0.260 | 0.305 | 0.345 | 0.345 | 0.240 | 0.355 | 0.250 | 0.325 | -45° |
| 45 | 0.300 | 0.310 | 0.300 | 0.350 | 0.260 | 0.310 | 0.345 | 0.345 | 0.245 | 0.355 | 0.250 | 0.325 | -45° |
| 46 | 0.310 | 0.310 | 0.305 | 0.355 | 0.270 | 0.310 | 0.355 | 0.350 | 0.250 | 0.360 | 0.260 | 0.330 | -45° |
| 47 | 0.295 | 0.320 | 0.295 | 0.360 | 0.260 | 0.320 | 0.345 | 0.355 | 0.240 | 0.360 | 0.250 | 0.335 | -45° |
| 48 | 0.290 | 0.315 | 0.290 | 0.355 | 0.255 | 0.315 | 0.340 | 0.350 | 0.235 | 0.365 | 0.245 | 0.335 | -45° |
| 49 | 0.300 | 0.320 | 0.295 | 0.360 | 0.265 | 0.315 | 0.345 | 0.355 | 0.245 | 0.365 | 0.255 | 0.335 | -45° |
| 50 | 0.305 | 0.315 | 0.300 | 0.350 | 0.265 | 0.315 | 0.345 | 0.345 | 0.245 | 0.360 | 0.255 | 0.330 | -45° |
| 51 | 0.305 | 0.320 | 0.295 | 0.355 | 0.260 | 0.315 | 0.350 | 0.350 | 0.240 | 0.360 | 0.250 | 0.335 | -45° |
| 52 | 0.305 | 0.320 | 0.300 | 0.365 | 0.265 | 0.320 | 0.350 | 0.355 | 0.250 | 0.365 | 0.255 | 0.340 | -45° |
| 53 | 0.300 | 0.320 | 0.295 | 0.355 | 0.260 | 0.315 | 0.345 | 0.350 | 0.245 | 0.360 | 0.255 | 0.330 | -45° |
| 54 | 0.290 | 0.320 | 0.285 | 0.355 | 0.255 | 0.315 | 0.340 | 0.350 | 0.235 | 0.360 | 0.245 | 0.330 | -45° |
| 55 | 0.295 | 0.315 | 0.295 | 0.350 | 0.260 | 0.315 | 0.345 | 0.345 | 0.245 | 0.360 | 0.250 | 0.330 | -45° |
| 56 | 0.300 | 0.325 | 0.295 | 0.360 | 0.260 | 0.315 | 0.345 | 0.350 | 0.245 | 0.360 | 0.255 | 0.335 | -45° |
| 57 | 0.300 | 0.315 | 0.295 | 0.355 | 0.260 | 0.310 | 0.340 | 0.345 | 0.240 | 0.355 | 0.250 | 0.330 | -45° |
| 58 | 0.295 | 0.325 | 0.290 | 0.365 | 0.255 | 0.320 | 0.340 | 0.355 | 0.240 | 0.365 | 0.250 | 0.340 | -45° |
| 59 | 0.300 | 0.320 | 0.295 | 0.360 | 0.265 | 0.320 | 0.345 | 0.350 | 0.245 | 0.365 | 0.255 | 0.335 | -45° |
| 60 | 0.285 | 0.325 | 0.285 | 0.360 | 0.255 | 0.320 | 0.340 | 0.355 | 0.240 | 0.365 | 0.250 | 0.335 | -45° |
| 61 | 0.300 | 0.320 | 0.295 | 0.355 | 0.260 | 0.315 | 0.340 | 0.350 | 0.240 | 0.355 | 0.250 | 0.330 | -45° |
| 62 | 0.300 | 0.325 | 0.295 | 0.360 | 0.260 | 0.320 | 0.340 | 0.355 | 0.240 | 0.365 | 0.250 | 0.335 | -45° |
| 63 | 0.300 | 0.315 | 0.300 | 0.355 | 0.260 | 0.310 | 0.350 | 0.350 | 0.250 | 0.360 | 0.255 | 0.330 | -45° |
| 64 | 0.305 | 0.320 | 0.300 | 0.360 | 0.265 | 0.315 | 0.350 | 0.350 | 0.245 | 0.365 | 0.260 | 0.335 | -45° |
| 65 | 0.305 | 0.315 | 0.300 | 0.355 | 0.265 | 0.315 | 0.345 | 0.345 | 0.245 | 0.360 | 0.255 | 0.330 | -45° |

| | | | | | | | | | | | | | |
|---|---|---|---|---|---|---|---|---|---|---|---|---|---|
| 66 | 0.300 | 0.320 | 0.300 | 0.355 | 0.265 | 0.310 | 0.345 | 0.350 | 0.245 | 0.360 | 0.255 | 0.325 | -45° |
| 67 | 0.300 | 0.320 | 0.295 | 0.355 | 0.265 | 0.320 | 0.350 | 0.350 | 0.250 | 0.360 | 0.260 | 0.335 | -45° |
| 68 | 0.300 | 0.320 | 0.295 | 0.355 | 0.265 | 0.315 | 0.345 | 0.350 | 0.240 | 0.360 | 0.255 | 0.330 | -45° |

**Supplementary Table 5. The coil placements of the clinical MDD cohort are shown in parameter space.**

| Sub ID | Treatment Group | Position ($s$) | | Orientation ($\theta$) |
|---|---|---|---|---|
| | | $p_{NZ}$ | $p_{AL}$ | |
| 1 | Left PFC | 0.300 | 0.285 | -45° |
| 2 | Left PFC | 0.220 | 0.435 | -45° |
| 3 | Left PFC | 0.300 | 0.345 | -45° |
| 4 | Left PFC | 0.120 | 0.480 | -45° |
| 5 | Left PFC | 0.155 | 0.250 | -45° |
| 6 | Left PFC | 0.215 | 0.435 | -45° |
| 7 | Left PFC | 0.270 | 0.280 | -45° |
| 8 | Left PFC | 0.295 | 0.360 | -45° |
| 9 | Left PFC | 0.345 | 0.380 | -45° |
| 10 | Left PFC | 0.410 | 0.365 | -45° |
| 11 | Left PFC | 0.380 | 0.365 | -45° |
| 12 | Left PFC | 0.365 | 0.335 | -45° |
| 13 | Left PFC | 0.345 | 0.350 | -45° |
| 14 | Left PFC | 0.315 | 0.350 | -45° |
| 15 | Left PFC | 0.380 | 0.345 | -45° |
| 16 | Left PFC | 0.420 | 0.280 | -45° |
| 17 | Left PFC | 0.355 | 0.310 | -45° |
| 18 | Left PFC | 0.385 | 0.355 | -45° |
| 19 | Left PFC | 0.415 | 0.315 | -45° |
| 20 | Left PFC | 0.325 | 0.325 | -45° |
| 21 | Left PFC | 0.270 | 0.295 | -45° |
| 22 | Left PFC | 0.330 | 0.340 | -45° |
| 23 | Left PFC | 0.325 | 0.295 | -45° |
| 24 | Left PFC | 0.420 | 0.385 | -45° |
| 25 | Left PFC | 0.385 | 0.370 | -45° |
| 26 | Left PFC | 0.350 | 0.315 | -45° |
| 27 | Left PFC | 0.370 | 0.320 | -45° |
| 28 | Right PFC | 0.325 | 0.540 | -45° |
| 29 | Right PFC | 0.360 | 0.555 | -45° |
| 30 | Right PFC | 0.325 | 0.580 | -45° |
| 31 | Right PFC | 0.295 | 0.745 | -45° |
| 32 | Right PFC | 0.300 | 0.640 | -45° |
| 33 | Right PFC | 0.180 | 0.725 | -45° |

**Supplementary Table 6. The coil placements of the clinical AVH cohort are shown in parameter space.**

| Sub ID | Treatment Group | Position ($s$) | | Orientation |
|---|---|---|---|---|
| | | $p_{NZ}$ | $p_{AL}$ | ($\theta$) |
| 1 | Active | 0.495 | 0.135 | -46.2° |
| 2 | Active | 0.785 | 0.170 | -2.8° |
| 3 | Active | 0.350 | 0.140 | -71.8° |
| 4 | Active | 0.710 | 0.190 | -13.7° |
| 5 | Active | 0.635 | 0.160 | -31.7° |
| 6 | Active | 0.740 | 0.140 | -17.2° |
| 7 | Active | 0.660 | 0.120 | -27.3° |
| 8 | Active | 0.615 | 0.150 | -33.9° |
| 9 | Active | 0.550 | 0.140 | -41° |
| 10 | Active | 0.610 | 0.210 | -36.3° |
| 11 | Active | 0.570 | 0.130 | -37.2° |
| 12 | Active | 0.685 | 0.135 | -25.6° |
| 13 | Active | 0.510 | 0.130 | -50.9° |
| 14 | Active | 0.565 | 0.825 | -40.1° |
| 15 | Active | 0.705 | 0.135 | -23.4° |

#### Supplementary Table 7. CPC positions of MDD simulation experiment.

| Pos ID | Position (*s*) | | Pos ID | Position (*s*) | | Pos ID | Position (*s*) | | Pos ID | Position (*s*) | |
|---|---|---|---|---|---|---|---|---|---|---|---|
| | $p_{NZ}$ | $p_{AL}$ | | $p_{NZ}$ | $p_{AL}$ | | $p_{NZ}$ | $p_{AL}$ | | $p_{NZ}$ | $p_{AL}$ |
| 1 | 0.077 | 0.373 | 36 | 0.164 | 0.439 | 71 | 0.239 | 0.319 | 106 | 0.319 | 0.413 |
| 2 | 0.081 | 0.413 | 37 | 0.167 | 0.376 | 72 | 0.241 | 0.426 | 107 | 0.321 | 0.349 |
| 3 | 0.084 | 0.347 | 38 | 0.169 | 0.476 | 73 | 0.244 | 0.361 | 108 | 0.323 | 0.451 |
| 4 | 0.090 | 0.389 | 39 | 0.170 | 0.306 | 74 | 0.246 | 0.463 | 109 | 0.326 | 0.389 |
| 5 | 0.093 | 0.321 | 40 | 0.171 | 0.414 | 75 | 0.247 | 0.291 | 110 | 0.329 | 0.321 |
| 6 | 0.094 | 0.427 | 41 | 0.176 | 0.350 | 76 | 0.249 | 0.401 | 111 | 0.331 | 0.427 |
| 7 | 0.099 | 0.363 | 42 | 0.177 | 0.453 | 77 | 0.251 | 0.336 | 112 | 0.334 | 0.364 |
| 8 | 0.100 | 0.464 | 43 | 0.179 | 0.277 | 78 | 0.254 | 0.440 | 113 | 0.336 | 0.466 |
| 9 | 0.103 | 0.403 | 44 | 0.180 | 0.390 | 79 | 0.257 | 0.377 | 114 | 0.339 | 0.404 |
| 10 | 0.106 | 0.337 | 45 | 0.183 | 0.323 | 80 | 0.259 | 0.477 | 115 | 0.343 | 0.339 |
| 11 | 0.109 | 0.441 | 46 | 0.186 | 0.429 | 81 | 0.260 | 0.309 | 116 | 0.344 | 0.443 |
| 12 | 0.111 | 0.379 | 47 | 0.189 | 0.366 | 82 | 0.263 | 0.417 | 117 | 0.347 | 0.380 |
| 13 | 0.113 | 0.479 | 48 | 0.190 | 0.467 | 83 | 0.266 | 0.351 | 118 | 0.353 | 0.419 |
| 14 | 0.114 | 0.310 | 49 | 0.191 | 0.296 | 84 | 0.267 | 0.454 | 119 | 0.356 | 0.354 |
| 15 | 0.116 | 0.419 | 50 | 0.193 | 0.406 | 85 | 0.269 | 0.280 | 120 | 0.357 | 0.457 |
| 16 | 0.120 | 0.353 | 51 | 0.196 | 0.340 | 86 | 0.270 | 0.393 | 121 | 0.360 | 0.394 |
| 17 | 0.121 | 0.456 | 52 | 0.199 | 0.444 | 87 | 0.273 | 0.326 | 122 | 0.366 | 0.433 |
| 18 | 0.124 | 0.394 | 53 | 0.201 | 0.381 | 88 | 0.276 | 0.431 | 123 | 0.369 | 0.370 |
| 19 | 0.127 | 0.327 | 54 | 0.203 | 0.481 | 89 | 0.279 | 0.367 | 124 | 0.374 | 0.410 |
| 20 | 0.130 | 0.433 | 55 | 0.204 | 0.313 | 90 | 0.280 | 0.469 | 125 | 0.381 | 0.386 |
| 21 | 0.133 | 0.369 | 56 | 0.206 | 0.420 | 91 | 0.281 | 0.299 | | | |
| 22 | 0.134 | 0.470 | 57 | 0.210 | 0.356 | 92 | 0.283 | 0.407 | | | |
| 23 | 0.136 | 0.300 | 58 | 0.211 | 0.459 | 93 | 0.287 | 0.341 | | | |
| 24 | 0.137 | 0.409 | 59 | 0.213 | 0.284 | 94 | 0.289 | 0.446 | | | |
| 25 | 0.140 | 0.343 | 60 | 0.214 | 0.396 | 95 | 0.291 | 0.383 | | | |
| 26 | 0.143 | 0.447 | 61 | 0.217 | 0.330 | 96 | 0.294 | 0.316 | | | |
| 27 | 0.144 | 0.270 | 62 | 0.220 | 0.434 | 97 | 0.297 | 0.423 | | | |
| 28 | 0.146 | 0.384 | 63 | 0.223 | 0.371 | 98 | 0.300 | 0.359 | | | |
| 29 | 0.149 | 0.317 | 64 | 0.224 | 0.473 | 99 | 0.301 | 0.460 | | | |
| 30 | 0.150 | 0.423 | 65 | 0.226 | 0.301 | 100 | 0.304 | 0.399 | | | |
| 31 | 0.154 | 0.360 | 66 | 0.227 | 0.411 | 101 | 0.307 | 0.331 | | | |
| 32 | 0.156 | 0.461 | 67 | 0.231 | 0.346 | 102 | 0.310 | 0.437 | | | |
| 33 | 0.157 | 0.289 | 68 | 0.233 | 0.449 | 103 | 0.313 | 0.374 | | | |
| 34 | 0.159 | 0.400 | 69 | 0.234 | 0.273 | 104 | 0.314 | 0.474 | | | |
| 35 | 0.161 | 0.333 | 70 | 0.236 | 0.387 | 105 | 0.316 | 0.304 | | | |

#### Supplementary Table 8. CPC positions of AVH simulation experiment.

| Pos ID | Position ($s$) | | Pos ID | Position ($s$) | | Pos ID | Position ($s$) | | Pos ID | Position ($s$) | |
|---|---|---|---|---|---|---|---|---|---|---|---|
| | $p_{NZ}$ | $p_{AL}$ | | $p_{NZ}$ | $p_{AL}$ | | $p_{NZ}$ | $p_{AL}$ | | $p_{NZ}$ | $p_{AL}$ |
| 1 | 0.086 | 0.093 | 36 | 0.516 | 0.146 | 71 | 0.670 | 0.074 | 106 | 0.757 | 0.217 |
| 2 | 0.120 | 0.110 | 37 | 0.524 | 0.080 | 72 | 0.671 | 0.267 | 107 | 0.759 | 0.344 |
| 3 | 0.134 | 0.147 | 38 | 0.537 | 0.124 | 73 | 0.674 | 0.173 | 108 | 0.760 | 0.083 |
| 4 | 0.141 | 0.081 | 39 | 0.550 | 0.157 | 74 | 0.677 | 0.314 | 109 | 0.761 | 0.270 |
| 5 | 0.154 | 0.126 | 40 | 0.559 | 0.099 | 75 | 0.680 | 0.236 | 110 | 0.764 | 0.177 |
| 6 | 0.169 | 0.159 | 41 | 0.563 | 0.186 | 76 | 0.683 | 0.121 | 111 | 0.767 | 0.317 |
| 7 | 0.176 | 0.100 | 42 | 0.571 | 0.139 | 77 | 0.684 | 0.286 | 112 | 0.770 | 0.239 |
| 8 | 0.190 | 0.139 | 43 | 0.576 | 0.210 | 78 | 0.689 | 0.199 | 113 | 0.773 | 0.127 |
| 9 | 0.197 | 0.066 | 44 | 0.580 | 0.064 | 79 | 0.690 | 0.331 | 114 | 0.774 | 0.289 |
| 10 | 0.211 | 0.117 | 45 | 0.584 | 0.169 | 80 | 0.693 | 0.256 | 115 | 0.779 | 0.203 |
| 11 | 0.224 | 0.151 | 46 | 0.590 | 0.233 | 81 | 0.696 | 0.156 | 116 | 0.783 | 0.259 |
| 12 | 0.231 | 0.089 | 47 | 0.593 | 0.116 | 82 | 0.699 | 0.304 | 117 | 0.786 | 0.160 |
| 13 | 0.246 | 0.131 | 48 | 0.597 | 0.196 | 83 | 0.701 | 0.221 | 118 | 0.794 | 0.101 |
| 14 | 0.267 | 0.107 | 49 | 0.606 | 0.150 | 84 | 0.703 | 0.347 | 119 | 0.807 | 0.140 |
| 15 | 0.280 | 0.144 | 50 | 0.611 | 0.219 | 85 | 0.704 | 0.094 | 120 | 0.816 | 0.069 |
| 16 | 0.287 | 0.076 | 51 | 0.614 | 0.087 | 86 | 0.706 | 0.274 | 121 | 0.829 | 0.117 |
| 17 | 0.301 | 0.123 | 52 | 0.619 | 0.180 | 87 | 0.709 | 0.183 | 122 | 0.841 | 0.153 |
| 18 | 0.314 | 0.156 | 53 | 0.624 | 0.240 | 88 | 0.711 | 0.321 | | | |
| 19 | 0.323 | 0.096 | 54 | 0.627 | 0.130 | 89 | 0.714 | 0.243 | | | |
| 20 | 0.336 | 0.136 | 55 | 0.629 | 0.290 | 90 | 0.717 | 0.136 | | | |
| 21 | 0.343 | 0.060 | 56 | 0.633 | 0.204 | 91 | 0.719 | 0.293 | | | |
| 22 | 0.357 | 0.113 | 57 | 0.634 | 0.336 | 92 | 0.723 | 0.209 | | | |
| 23 | 0.370 | 0.149 | 58 | 0.637 | 0.260 | 93 | 0.724 | 0.337 | | | |
| 24 | 0.379 | 0.084 | 59 | 0.640 | 0.161 | 94 | 0.726 | 0.057 | | | |
| 25 | 0.391 | 0.127 | 60 | 0.643 | 0.309 | 95 | 0.727 | 0.263 | | | |
| 26 | 0.404 | 0.160 | 61 | 0.646 | 0.227 | 96 | 0.730 | 0.166 | | | |
| 27 | 0.413 | 0.103 | 62 | 0.649 | 0.106 | 97 | 0.733 | 0.311 | | | |
| 28 | 0.426 | 0.141 | 63 | 0.650 | 0.280 | 98 | 0.736 | 0.230 | | | |
| 29 | 0.434 | 0.070 | 64 | 0.653 | 0.189 | 99 | 0.739 | 0.111 | | | |
| 30 | 0.447 | 0.119 | 65 | 0.656 | 0.326 | 100 | 0.740 | 0.281 | | | |
| 31 | 0.460 | 0.153 | 66 | 0.659 | 0.249 | 101 | 0.744 | 0.193 | | | |
| 32 | 0.469 | 0.091 | 67 | 0.661 | 0.143 | 102 | 0.746 | 0.327 | | | |
| 33 | 0.481 | 0.133 | 68 | 0.663 | 0.297 | 103 | 0.749 | 0.251 | | | |
| 34 | 0.490 | 0.053 | 69 | 0.667 | 0.213 | 104 | 0.751 | 0.147 | | | |
| 35 | 0.503 | 0.110 | 70 | 0.669 | 0.341 | 105 | 0.754 | 0.300 | | | |

**Supplementary Table 9. The optimal TMS coil placements of MDD simulation experiment.**

| Sub ID | Position ($s$) | | Orientation ($\theta$) |
|---|---|---|---|
| | $p_{NZ}$ | $p_{AL}$ | |
| 1 | 0.336 | 0.306 | -45° |
| 2 | 0.201 | 0.381 | -120° |
| 3 | 0.189 | 0.366 | -30° |
| 4 | 0.167 | 0.376 | -15° |
| 5 | 0.321 | 0.299 | -60° |
| 6 | 0.189 | 0.366 | -30° |
| 7 | 0.249 | 0.401 | -120° |
| 8 | 0.201 | 0.381 | -105° |
| 9 | 0.189 | 0.366 | -90° |
| 10 | 0.210 | 0.356 | -45° |
| 11 | 0.210 | 0.356 | -60° |
| 12 | 0.210 | 0.356 | -60° |
| 13 | 0.316 | 0.304 | -15° |
| 14 | 0.299 | 0.290 | -45° |
| 15 | 0.180 | 0.390 | -30° |
| 16 | 0.214 | 0.396 | -60° |
| 17 | 0.189 | 0.366 | -45° |
| 18 | 0.167 | 0.376 | -30° |
| 19 | 0.210 | 0.356 | -15° |
| 20 | 0.210 | 0.356 | -165° |
| 21 | 0.210 | 0.356 | -105° |
| 22 | 0.189 | 0.366 | 0° |
| 23 | 0.321 | 0.299 | -30° |
| 24 | 0.231 | 0.346 | -105° |
| 25 | 0.210 | 0.356 | -90° |
| 26 | 0.210 | 0.356 | -165° |
| 27 | 0.189 | 0.366 | -150° |

**Supplementary Table 10. The optimal TMS coil placements of AVH simulation experiment.**

| Sub ID | Position ($s$) | | Orientation ($\theta$) |
|---|---|---|---|
| | $p_{NZ}$ | $p_{AL}$ | |
| 1 | 0.436 | 0.159 | -135° |
| 2 | 0.401 | 0.159 | -135° |
| 3 | 0.654 | 0.347 | -150° |
| 4 | 0.344 | 0.159 | -150° |
| 5 | 0.626 | 0.324 | -150° |
| 6 | 0.401 | 0.159 | -165° |
| 7 | 0.626 | 0.323 | -135° |
| 8 | 0.633 | 0.336 | -150° |
| 9 | 0.401 | 0.159 | -135° |
| 10 | 0.373 | 0.159 | -150° |
| 11 | 0.314 | 0.156 | -135° |
| 12 | 0.154 | 0.126 | -45° |
| 13 | 0.374 | 0.159 | -165° |
| 14 | 0.403 | 0.159 | -135° |
| 15 | 0.401 | 0.159 | -135° |

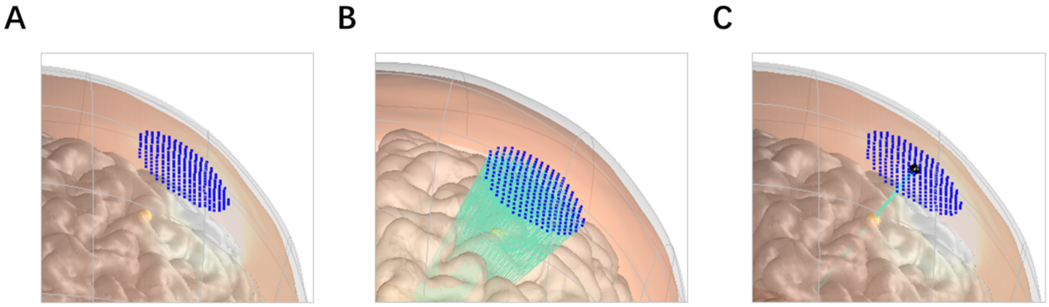

**Supplementary Figure 1. Steps of finding a scalp position based on a DLPFC site. (A)** For one DLPFC site (yellow sphere), we first found close by CPC positions (blue dots) within 30 mm of the site. **(B)** The scalp-normal vectors of these CPC positions were pointed to the cortical surface (cyan lines). Then we calculated the distance between the DLPFC site and the normal vectors. **(C)** Finally, we selected a CPC position (black sphere) in which the normal vector (cyan line) was closest to the DLPFC site (yellow sphere).

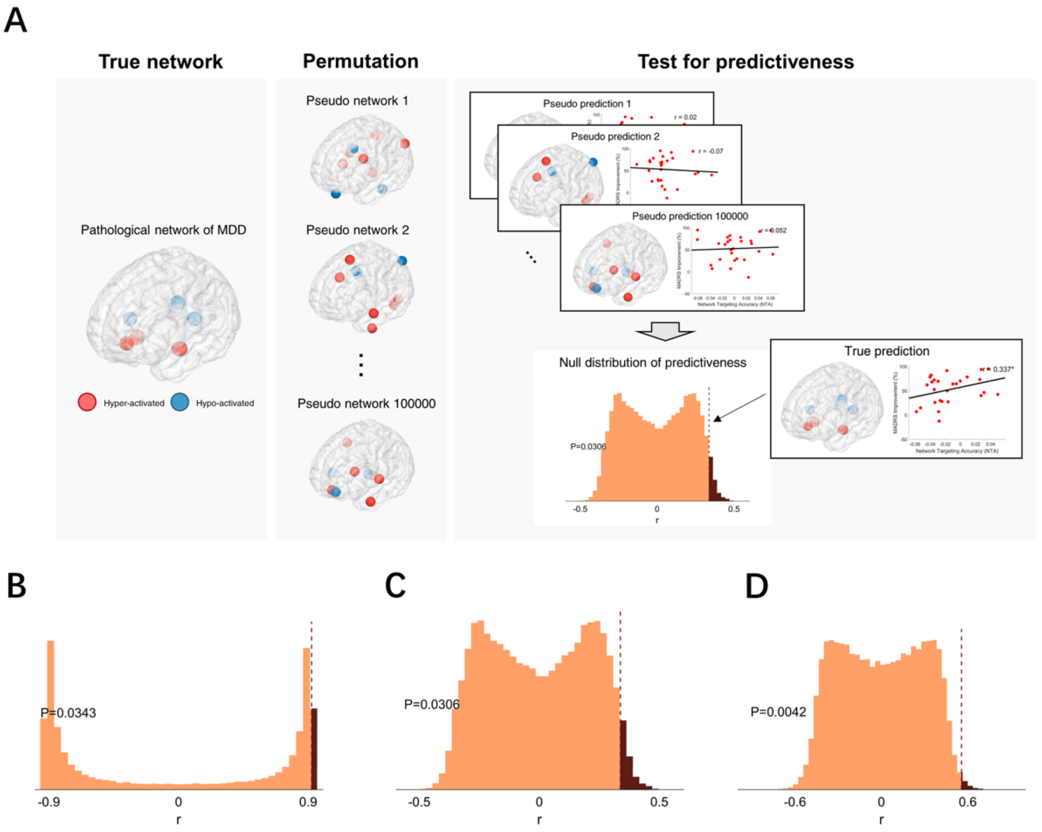


**Supplementary Figure 2. Permutation tests of targeting pseudo networks for assessing the predictiveness of NTA. (A)** Comparison between targeting the true pathological network and targeting pseudo networks. Pseudo networks generated by permutating the spatial distribution of hyper- and hypo-activation foci (Gray et al., 2020) were used as alternative networks of targets. NTAs based on the pseudo networks were used to predict the treatment efficacies on an illustrative cohort. For the cohort of patients, the permutation was conducted $10^5$ times to generate the null distribution and the ranked percentile on this distribution indicated the significant level of predictiveness of NTA. For a given pathological network (the first column), positions of the hypo- and hyper-activated foci were randomized around the cortex. The resultant pseudo networks (the second column) were used to implement NTA models to predict the site-wise treatment efficacies. Frequencies of each r-value of the pseudo predictions were accumulated as the null distribution. The ranked percentile in the null distribution determined the level of significance of the true prediction, i.e., predicting the treatment efficacy with NTA based on the pathological network (the third column). **(B)** The permutation test on continuous DLPFC sites with equation-based clinical efficacy ($N = 68$). **(C)** The permutation test on the clinical MDD cohort ($N = 27$). **(D)** The permutation test on the clinical AVH cohort ($N = 15$).

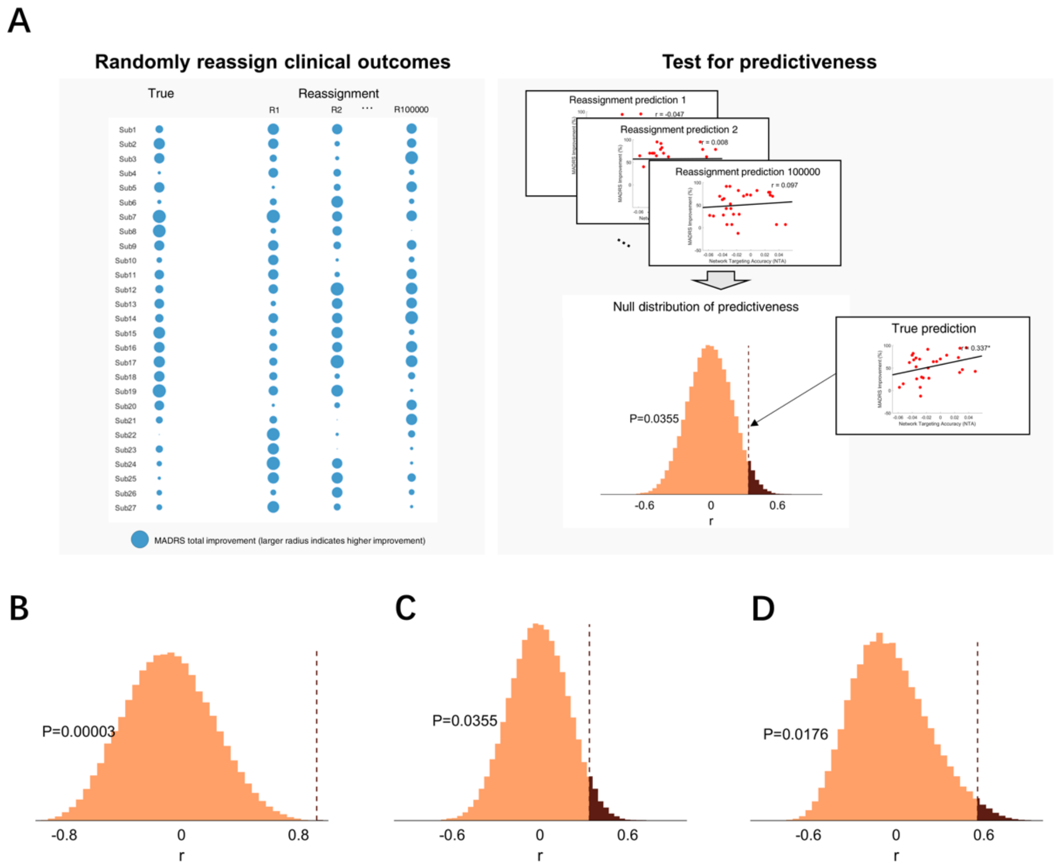


**Supplementary Figure 3. Permutation tests of randomized prediction for assessing the predictiveness of NTA. (A)** Comparison between true prediction and random prediction. We fixed the true pathological network and reassigned the clinical outcomes. NTA was used to predict the randomized treatment efficacies on an illustrative cohort. For the cohort of patients, the permutation was conducted $10^5$ times to generate the null distribution and the ranked percentile on this distribution indicated the significant level of predictiveness of NTA on true clinical outcomes. Each patient's clinical outcome was randomly reassigned to a different patient (the first column). Frequencies of each r-value from the randomized predictions were accumulated as the null distribution. The ranked percentile in the null distribution determined the level of significance of the true prediction, i.e., predicting the treatment efficacy with NTA based on the true labels (the second column). **(B)** The permutation test on empirical

DLPFC sites of a large depression cohort (*N* = 68). **(C)** The permutation test on the clinical cohort of MDD (*N* = 27). **(D)** The permutation test on the clinical AVH cohort (*N* = 15).

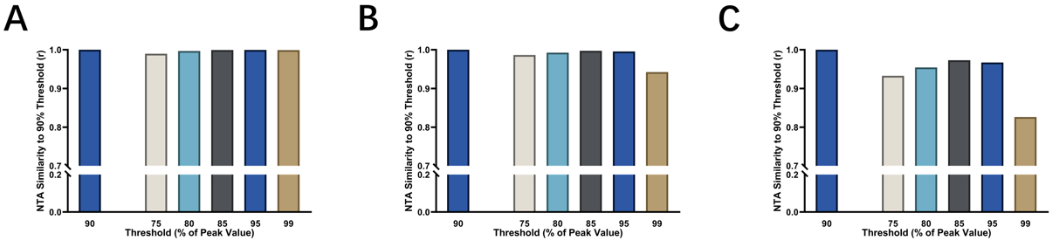


**Supplementary Figure 4. Robustness of E-field threshold. (A)** Correlation between NTA from using the current 90% threshold and NTA from using other thresholds, 75%, 80%, 85%, 95%, and 99%, at the empirical DLPFC sites based on a large depression cohort ($N = 68$). **(B)** Correlation on the clinical MDD cohort ($N = 27$). **(C)** Correlation on the clinical AVH cohort ($N = 15$).

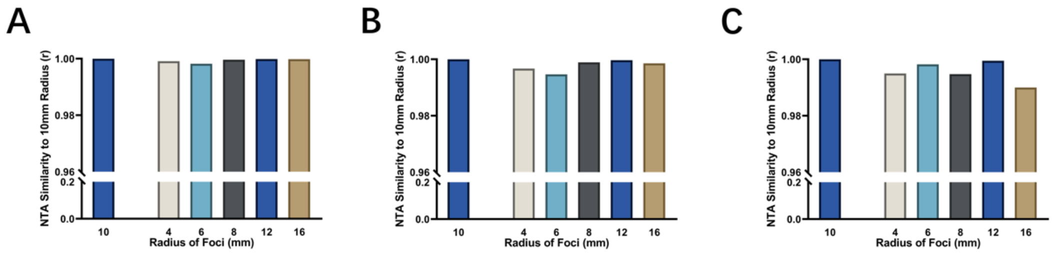


**Supplementary Figure 5. Robustness of radii of foci of the pathological network. (A)** Correlation between NTA from using the current 10 mm radius and NTA from using other radii, included 4 mm, 6 mm, 8 mm, 12 mm, and 16 mm, on the empirical DLPFC sites based on a large depression cohort (*N* = 68). **(B)** Correlation on the clinical MDD cohort (*N* = 27). **(C)** Correlation on the clinical AVH cohort (*N* = 15).

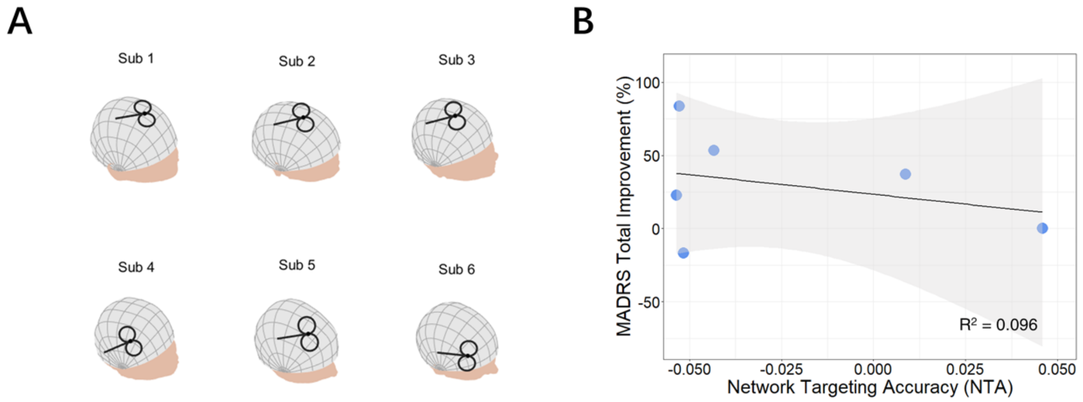


**Supplementary Figure 6. Network targeting accuracy failed to predict treatment efficacy in the clinical MDD cohort who received TMS in the right PFC. (A)** Coil placements of the right PFC patients are shown on individual head models. **(B)** Correlation between the NTA and MADRS total improvement ($N = 6$, $p = 0.725$, one-tailed).

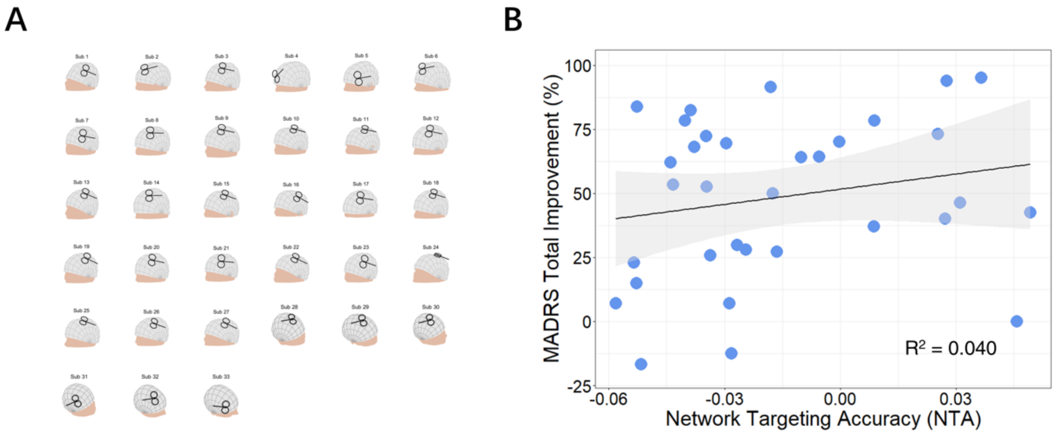


**Supplementary Figure 7. Network targeting accuracy failed to predict treatment efficacy in the clinical MDD cohort that received TMS on both sides of PFC. (A)** Coil placements of the double-side PFC patients shown on individual head models. **(B)** Correlation between the NTA and MADRS total improvement ($N = 33$, $p = 0.131$, one-tailed).

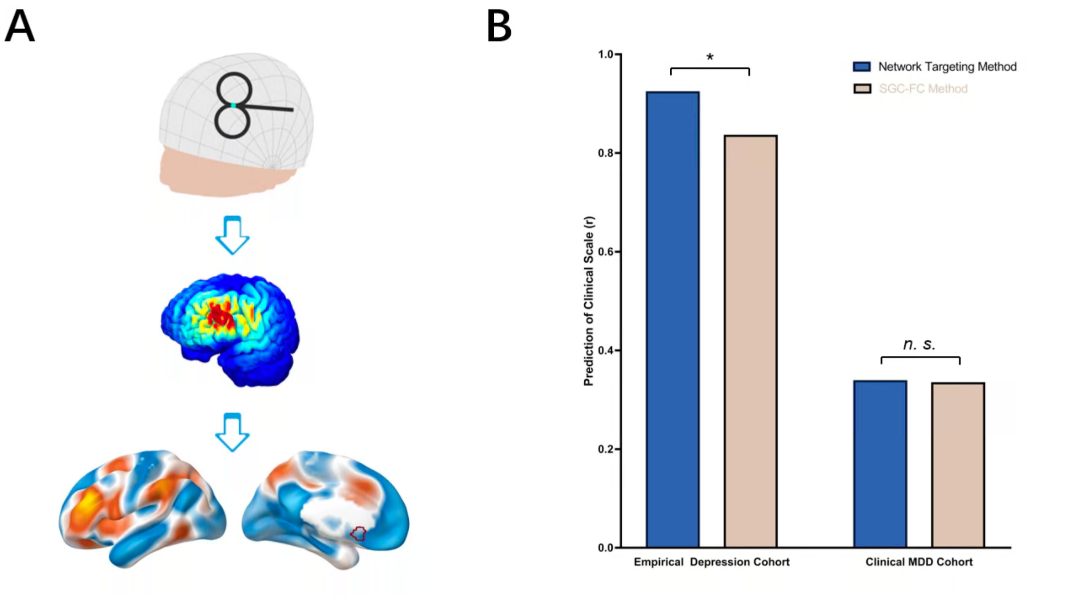


**Supplementary Figure 8. Comparing network targeting with the SGC-FC method in depression cohorts. (A)** The pipeline of the SGC-FC method (Fox et al., 2012). Instead of using a sphere to represent the cortical TMS effect, we used the E-field and calculated the 'Stimulation Network' with the same coil sets as for our network targeting method. The SGC ROI on the gray matter was constructed by placing a 10 mm radius sphere at MNI coordinates [6, 16, -10] (Fox et al., 2013). The targeting score of the SGC-FC method was calculated by averaging the sign-reversed functional connectivity within the SGC ROI. **(B)** The comparison between network targeting and the SGC-FC method in MDD cohorts. For the SGC-FC method, the targeting score predicted HDRS improvement in the empirical DLPFC sites based on a large depression cohort ($N$ = 68, r = 0.835, $p$ = $3.64\times10^{-4}$, one-tailed) and predicted MADRS improvement in the clinical MDD cohort ($N$ = 27, r = 0.333, $p$ = 0.048, one-tailed). Network targeting method had slightly higher r than the SGC-FC method in empirical cohort (t = 2.149, $p$ = 0.030, one-tailed) and similar to the SGC-FC method in clinical cohort (t = 0.032, $p$ = 0.487, one-tailed).

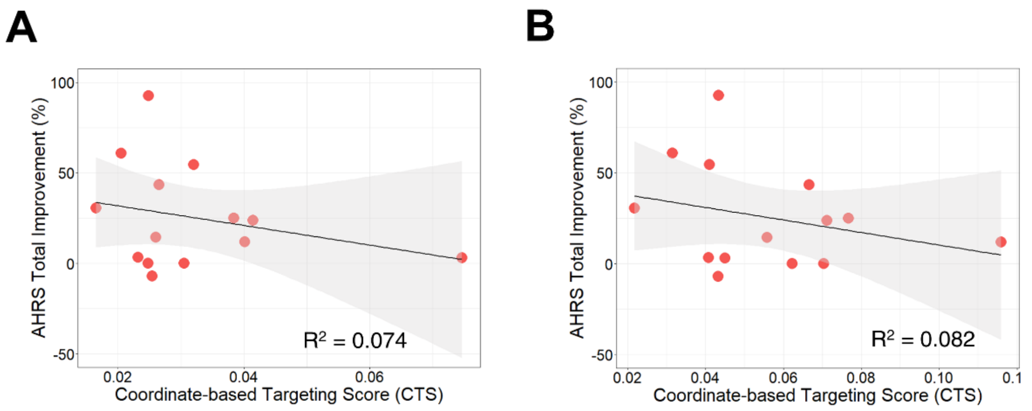


**Supplementary Figure 9. Coordinates-based targeting method in AVH treatment. (A)** TP3 (Herwig et al., 2003) failed in predicting AHRS total improvement ($N$ = 14, r = -0.272, p = 0.827, one-tailed). **(B)** L. Wernicke (Hoffman et al., 2013) failed in predicting AHRS total improvement ($N$ = 14, r = -0.285, p = 0.839, one-tailed).
